# Electron confurcation drives photosynthetic H_2_ production in cyanobacteria

**DOI:** 10.64898/2026.06.25.734504

**Authors:** Gyana Prakash Mahapatra, Nadine Strabel, Christian Lorent, Anuj Kumar, Stefan Bohn, Max A. Klamke, Marko Böhm, Christian Teutloff, Sven T. Stripp, Ingo Zebger, Jens Appel, Jan M. Schuller, Kirstin Gutekunst

**Affiliations:** Philipps-University Marburg, Department of Chemistry and Center for Synthetic Microbiology (SYNMIKRO), Karl-von-Frisch-Strasse 14, 35043 Marburg, Germany; Universität Kassel, Institut für Biologie, Heinrich-Plett-Straße 40, 34132 Kassel, Germany; Institut für Chemie, PC14, Physikalische und Biophysikalische Chemie, Technische Universität Berlin, Straße des 17. Juni 135, 10623 Berlin, Germany; Institute of Structural Biology, Helmholtz Munich, Ingolstädter Landstraße, 85764c Neuherberg, Germany; Freie Universität Berlin, Fachbereich Physik, Arnimallee 14, 14195 Berlin, Germany; Spectroscopy & Biocatalysis, Institute of Chemistry, University of Potsdam, Karl-Liebknecht-Strasse 24/25, 14476 Potsdam, Germany; Microbes-for-Climate (M4C) Cluster of Excellence, Marburg, Germany

## Abstract

Cyanobacteria are major contributors to global photosynthesis and are intensively studied for sustainable green H_2_ production. Central to this process is the bidirectional [NiFe]-hydrogenase HoxEFUYH, yet its physiological redox partners have remained unresolved. Ferredoxin, NAD(H), and NADP(H) have been proposed as partners, but the lack of active enzyme preparations has prevented a definitive assignment. Here, we purified the intact HoxEFUYH complex from *Synechocystis* sp. PCC 6803 under strictly anaerobic conditions and reveal its function as both a bifurcating and confurcating hydrogenase. During H_2_ uptake, HoxEFUYH utilizes NAD^+^ and oxidized ferredoxin, whereas H_2_ production strictly requires both NADH and reduced ferredoxin; NADPH does not support either reaction. Combining high-resolution cryo-electron microscopy with biochemical and spectroscopic analyses, our data reveal that an flavin-containing reductase module is electronically connected to the catalytic [NiFe]-hydrogenase core through an extended chain of iron-sulfur clusters, defining the structural basis for bifurcating and confurcating electron flow. These findings fundamentally revise the physiological role of HoxEFUYH by showing that photosynthetic H_2_ production does not rely solely on photosynthetic electrons but instead couples reduced ferredoxin from the light reaction with NADH derived from “dark” carbohydrate oxidation. This requires reassessment of current strategies for green H_2_ production in cyanobacteria.

## Introduction

Cyanobacteria have long attracted attention as platforms for sustainable dihydrogen (H_2_) production because they combine photosynthesis with enzymatic hydrogen metabolism^1,2^. Central to this capability is the bidirectional [NiFe]-hydrogenase Hox, a soluble, cytosolic enzyme that catalyzes both H_2_ evolution and oxidation, linking cellular redox metabolism to molecular hydrogen^3–6^. Unlike strictly H_2_-evolving or H_2_-oxidizing hydrogenases, the cyanobacterial Hox system operates reversibly and can integrate electrons from multiple metabolic sources. Hydrogenases are generally classified into [NiFe]-, [FeFe]-, and [Fe]-hydrogenases based on the metal composition of their catalytic cofactors. Among these, [NiFe]-hydrogenases are typically associated with H_2_ metabolism and diverse physiological functions ranging from energy conservation to the balancing of redox reactions^7–9^.

In many cyanobacteria, including *Synechocystis* sp. PCC 6803, the [NiFe]-hydrogenase is encoded by the *hoxEFUYH* operon and assembles into a large multisubunit complex. Its catalytic core (HoxYH) harbors the Ni-Fe active site, while the associated reductase (diaphorase) module (HoxEFU) contains multiple iron-sulfur (FeS) clusters and one flavin mononucleotide (FMN) active site and mediates electron transfer (ET) with physiological redox partners. Nevertheless, the identity of these redox partners has been debated for decades. NAD(H), NADP(H), and ferredoxin (Fdx)_red/ox_ have been tested as potential electron carriers; however, due to methodological challenges and conflicting results, they have not yet been definitively identified. *In vitro* studies were conducted either on cell homogenates or on isolated hydrogenases^10–18^. Cell homogenates are not free of naturally occurring electron carriers and also contain, for example, transhydrogenases that can transfer electrons between redox partners, thereby complicating the unambiguous identification of the hydrogenase’s redox partners. *In vitro* tests with isolated hydrogenase were either unsuccessful or lacked clear evidence that the entire HoxEFUYH complex remained intact^19^. This is particularly critical because artificial electron donors (e.g., reduced methyl viologen or dithionite) can reduce subfractions of the Hox complex in the absence of the reductase module and lead to H_2_ production. In this sense, the addition of natural redox partners to non-intact HoxEFUYH complexes can also lead to false-positive results. Since it has not yet been accomplished to isolate the entire intact HoxEFUYH in its active form, all investigations to date on the quest of redox partners of cyanobacterial HoxEFUYH should therefore be interpreted with caution.

This uncertainty directly affects current models of the physiological role of HoxEFUYH. Based on current knowledge, it is assumed that HoxEFUYH accepts electrons from NADH, NADPH, or reduced ferredoxin for fermentative H_2_ production in the dark to balance the redox state of the cells. The photosynthetic light reaction and its main electron acceptor, the Calvin-Benson-Bassham (CBB) cycle are inactive in the dark. Illumination of anoxic dark-adapted cultures rapidly activates the photosynthetic electron transfer chain, whereas its major electron sink, the CBB cycle, becomes active only after a short delay^20,21^. This leads to a brief burst of so-called photosynthetic hydrogen production (photoH_2_) in which photoH_2_ acts as an electron buffer and the hydrogenase as an overflow valve^21^. From a biotechnological perspective, this transient light-driven photoH_2_ production is particularly attractive, as light energy is directly stored in the form of molecular hydrogen. Subsequently, the produced photoH_2_ is reoxidized, and as soon as O_2_ accumulates due to PSII activity, HoxEFUYH is inactivated^21^. PhotoH_2_ is currently believed to be either based on Fdx_red_ or NADPH delivered by PSI, while H_2_ uptake either transfers electrons onto NAD^+^ or alternatively possibly via the NDH-1 complex into the PQ pool of the photosynthetic electron transfer chain^16,17,21,22^.

Furthermore, these unexplained physiological observations also raise questions about the mechanism by which HoxEFUYH distributes electrons across different redox pathways. Notably, the cyanobacterial HoxEFU module exhibits structural similarity to the HydBC modules of bifurcating hydrogenases from acetogens, indicating that Hox may channel electron flow in a comparable manner^23– 26^. This possible mechanistic parallel is also important in the context of photobiological H_2_ production. From an applied perspective, the Hox system represents a promising entry point for photobiological H_2_ production^27,28^. In cyanobacteria, photosystem I (PSI) generates reduced ferredoxin using sunlight, which can, in principle, be directly funneled to hydrogenase-driven H_2_ evolution. Proof-of-concept studies, including engineered PSI–hydrogenase chimeras, have demonstrated that such coupling supports light-driven hydrogen production *in vitro* and *in vivo*^*2,29–33*^. These strategies highlight the potential of redirecting photosynthetic electrons away from native metabolic sinks towards H_2_ formation, converting solar energy into a clean, storable fuel.

Here, we show for the first time the isolation of the entire, active HoxEFUYH complex from *Synechocystis* sp. PCC 6803 in sufficient yield to perform biochemical, structural, and spectroscopic analysis. Using anaerobic purification, we obtained a homogeneous preparation that retains full H_2_-evolving and H_2_-oxidizing activity, enabling precise characterization of electron donor and acceptor usage and revealing mechanistic insights. Contrary to previous assumptions HoxEFUYH is a bifurcating and confurcating enzyme that utilizes the soluble electron carriers NAD(H) and ferredoxin_ox/red_. Accordingly, photoH_2_ production relies on a combination of electrons derived from NADH and Fdx_red_. This requires a reassessment of photoH_2_, as its production evidently involves the combination of electrons from carbohydrate oxidation (NADH) and the light reaction of photosynthesis (Fdx_red_). Our work establishes a mechanistic framework for understanding cyanobacterial H_2_ metabolism and provides the foundation for the rational engineering of hydrogenases and future photobiological H_2_ production platforms.

## Results

### Purification and bidirectional activity of the cyanobacterial [NiFe]-hydrogenase

To enable biochemical characterization, the hoxEFUYH operon from *Synechocystis* sp. PCC 6803 was engineered with an N-terminal Twin-Strep tag on the large hydrogenase subunit HoxH **(Fig. 1a)**. During purification, the enzyme was maintained under anaerobic and mildly reducing conditions (2 mM dithiothreitol, DTT). Size-exclusion chromatography (SEC) yielded a dominant, homogeneous high-molecular-weight species **(Extended Data Fig. 1a)**, and SDS-PAGE (sodium dodecyl sulfate polyacrylamide gel electrophoresis) together with mass spectrometry confirmed the presence of all five subunits **(Fig. 1a)**, establishing a robust basis for functional assays, high-resolution cryogenic electron microscopy (cryo-EM), and spectroscopic studies.

**Fig. 1:**
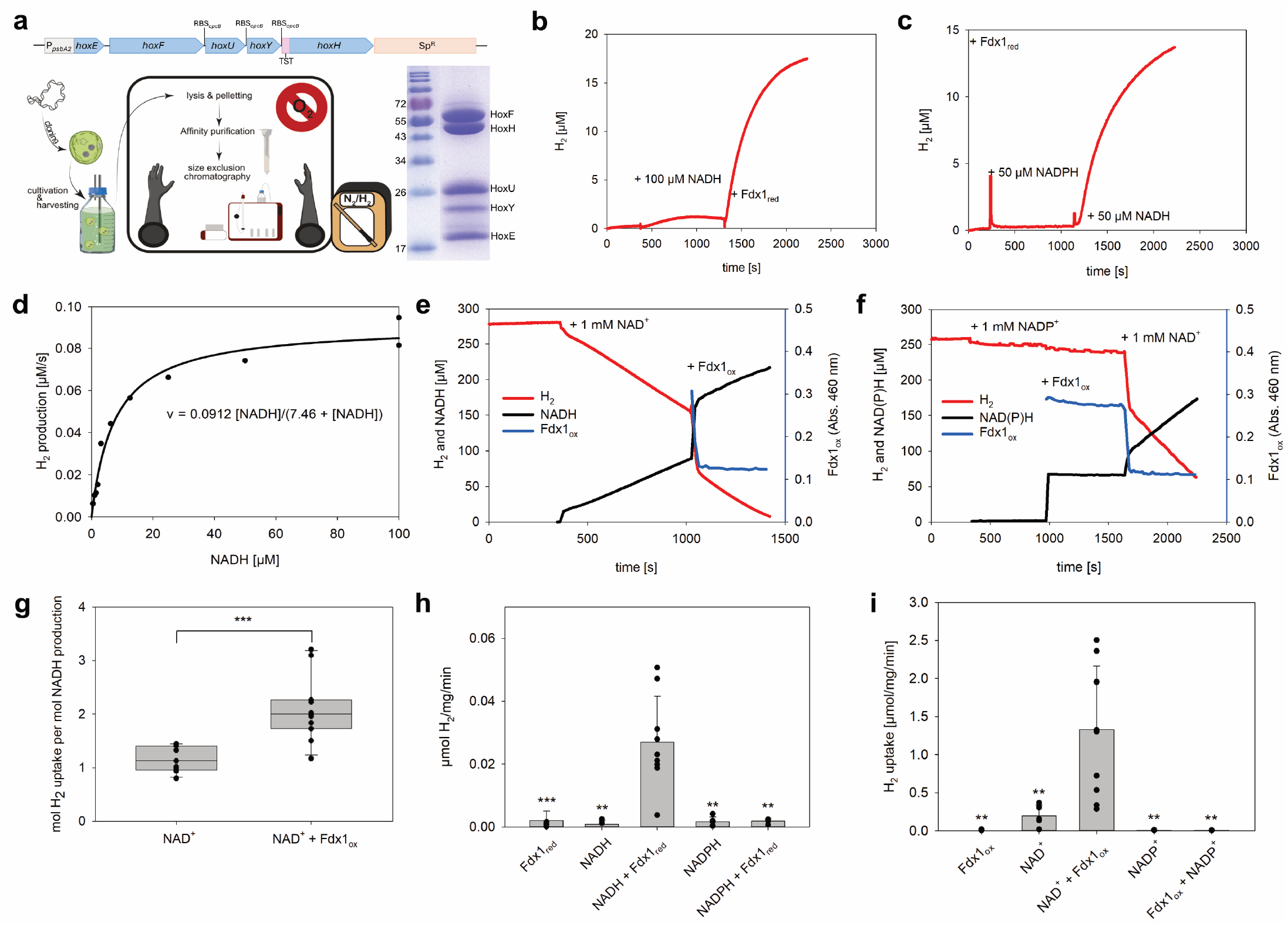
Purification and functional characterization of the cyanobacterial bidirectional [NiFe]-hydrogenase. **a**, Genetic organization of the engineered hoxEFUYH operon from Synechocystis sp. PCC 6803 and purification of the HoxEFUYH complex by size-exclusion chromatography and SDS-PAGE analysis confirms the presence of all five subunits. **b-d**, Hydrogen evolution activity of the purified enzyme with different electron donors. **b**, Addition of NADH alone does not result in detectable H_2_ production, whereas H_2_ evolution is observed upon addition of reduced ferredoxin. **c**, In the presence of reduced ferredoxin, NADH supports H_2_ production, while NADPH does not. **d**, Dependence of the H_2_ production rate on NADH concentration. **e-g**, Hydrogen oxidation activity of the purified enzyme. **e**, In the presence of NAD^+^, H_2_ is oxidized concomitantly with NADH formation; subsequent addition of oxidized ferredoxin markedly increases the rates of both processes. **f**, In the presence of oxidized ferredoxin and NADP^+^, no H_2_ oxidation is detected. **g**, Addition of NAD^+^ restores H_2_ oxidation activity. **h-i**, The [NiFe] hydrogenase HoxEFUYH of Synechocystis operates both in a confurcating mode with NADH and Fdxred for H_2_ evolution and a bifurcating mode with NAD^+^ and Fdxox during H_2_ uptake.

Under strictly anaerobic conditions, the purified HoxEFUYH complex was first tested for its capacity to catalyze H_2_ evolution. To maintain anaerobic conditions during the measurements, all assays contained an O_2_-scavenging system consisting of glucose, glucose oxidase, and catalase. Reduced ferredoxin was generated in situ using pyruvate:ferredoxin oxidoreductase (PFOR)^34^. Addition of NADH, NADPH or Fdx_red_ alone resulted in negligible H_2_ production at baseline levels **(Fig. 1b, c)**. In contrast, rapid H_2_ evolution was observed only when NADH was supplied together with Fdx_red_, demonstrating that efficient H_2_ production strictly requires the simultaneous presence of both electron donors **(Fig. 1b)**. The apparent K_M_ for NADH during confurcating H_2_ evolution in combination with Fdx_red_ was determined to be 7.48 µM **(Fig. 1d)**.

We next examined the reverse reaction, H_2_ oxidation. No H_2_ uptake was observed when NADP^+^ or Fdx_ox_ were added either alone or in combination **(Fig. 1f)**. In the presence of NAD^+^, slow H_2_ oxidation accompanied by NADH formation was detected **(Fig. 1e)**. Rapid H_2_ uptake was observed when NAD^+^ was added together with Fdx_ox_ **(Fig. 1e)**, indicating cooperative electron acceptance. Notably, the addition of exogenous FMN (100 µM) further increased H_2_ uptake rates under these conditions **(Extended Data Fig. 1d)**, supporting a role for FMN as a redox-active coenzyme in HoxEFUYH. This observation is consistent with previous studies showing that supplementation with exogenous FMN enhances electron-bifurcation activity in related hydrogenases^23,24,35^. Stoichiometric analysis revealed a 1:1 coupling between H_2_ oxidation and NADH formation when NAD^+^ was the sole electron acceptor, whereas the presence of Fdx_ox_ shifted the stoichiometry to approximately 2:1, consistent with simultaneous reduction of NAD^+^ and Fdx_ox_ **(Fig. 1g)**. Together, these data unambiguously establish that the HoxEFUYH complex is a bona fide confurcating [NiFe]-hydrogenase for H_2_ production with NADH and Fdx_red_ and a bifurcating [NiFe]-hydrogenase for H_2_ uptake with NAD^+^ and Fdx_ox_, while H_2_ uptake was also possible with NAD^+^ alone at significantly lower rates **(Fig. 1h, i)**.

### Overall structural organization and oligomerization of the HoxEFUYH complex

To determine the structural organization of the cyanobacterial bidirectional hydrogenase and provide a structural basis for the bifurcation and/or confurcation activities, we subjected the HoxEFUYH complex to cryo-EM SPA. All sample handling and grid preparation were performed anaerobically under strictly anaerobic reducing conditions (95% N_2_ and 5% H_2_) to preserve the native state of the enzyme. Analysis of the resulting cryo-EM dataset revealed a predominant dimeric population together with higher-order oligomeric assemblies **(Extended Data Fig. 2b)**. While these higher-order species were clearly visible in the dataset, pronounced preferred orientation prevented their reliable 3D reconstruction. In contrast, the dominant dimeric population yielded a reconstruction at an overall resolution of 2.30 Å **(Extended Data Fig. 2)**, allowing detailed structural analysis of the HoxEFUYH complex.

The structure reveals a dimeric heteropentameric assembly, hereafter referred to as HoxEFUYH_2_, exhibiting two-fold rotational symmetry (C2) **(Fig. 2a, b)**. At the protomer level, HoxEFUYH adopts an elongated architecture composed of five subunits that partition into two structurally and functionally distinct modules. Each protomer consists of the reductase module formed by HoxE, HoxF, and HoxU and the catalytic hydrogenase subunits HoxH and HoxY. The quality of the reconstruction enabled unambiguous modelling of all five subunits, multiple FeS clusters and the catalytic [NiFe] active site, indicating that the enzyme remained structurally intact throughout purification and vitrification. However, no density corresponding to bound FMN, NAD(H), indicating that this reconstruction represents a substrate and coenzyme-free (apo) state of HoxEFUYH. Notably, the complete complement of FeS clusters and the catalytic [NiFe] active site remained fully occupied.

**Fig. 2:**
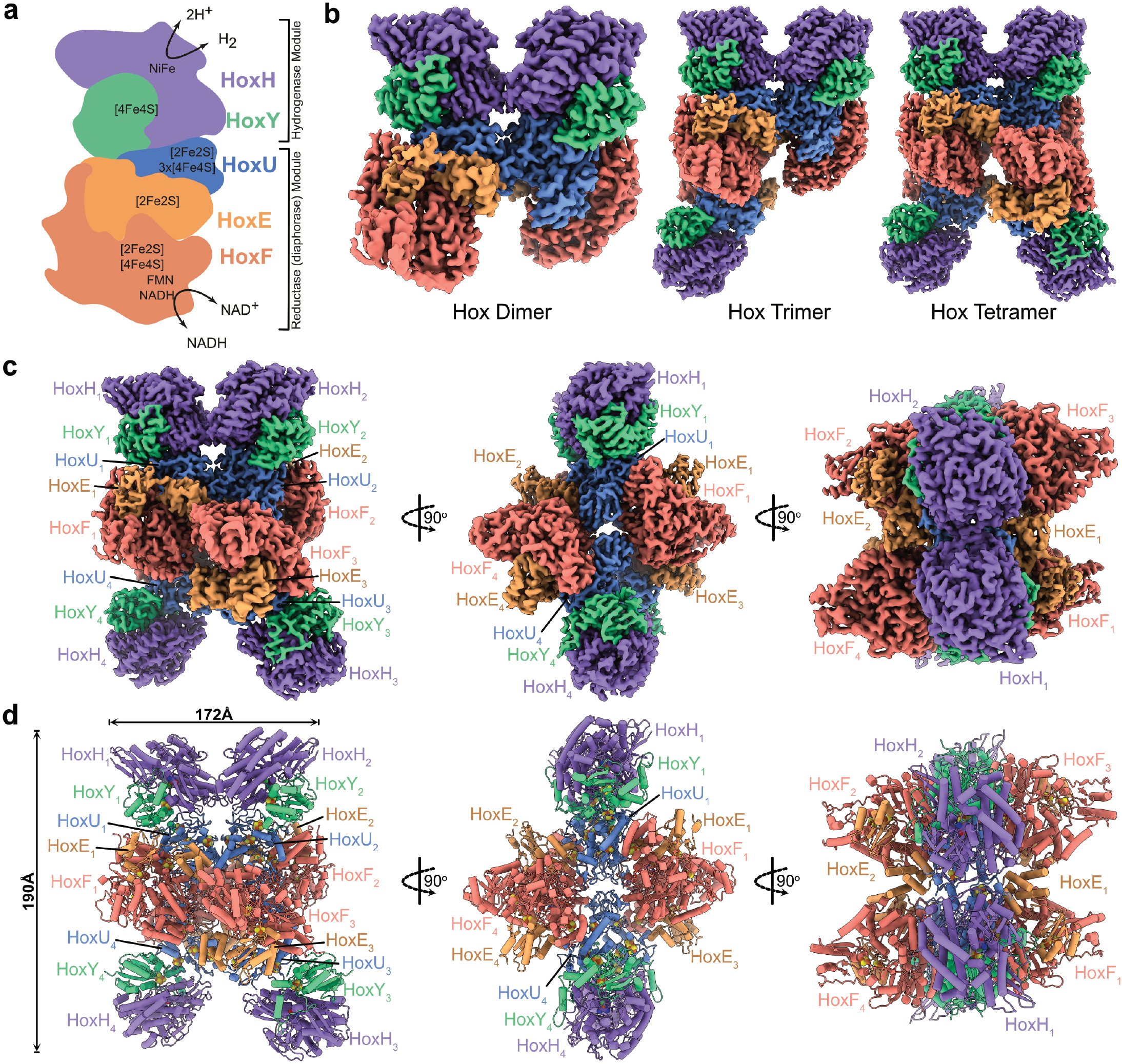
Structural organization and oligomeric assemblies of the cyanobacterial HoxEFUYH hydrogenase. **a**, Overall organization of the HoxEFUYH complex and distribution of redox cofactors within the hydrogenase and reductase (diaphorase) modules. **b**, Cryo-EM density maps of the different oligomeric assemblies identified in the holo dataset, including dimeric, trimeric, and tetrameric states. Individual subunits are colored consistently throughout. **c**, Orthogonal views of the tetrameric HoxEFUYH assembly shown as cryo-EM density maps. Individual protomers and subunits are labeled to highlight the spatial organization and intermolecular interfaces within the higher-order assembly. **d**, Atomic model shown in cartoon in the corresponding orientations as in panel c.

To improve angular sampling and capture catalytic states of the enzyme, cryo-EM grids were prepared anaerobically in the presence of FMN, NADH, and the zwitterionic detergent 3-([3-Cholamidopropyl]dimethylammonio)-2-hydroxy-1-propanesulfonate (CHAPSO). Under these conditions, tetrameric (HoxEFUYH_4_), trimeric (HoxEFUYH_3_), and dimeric (HoxEFUYH_2_) assemblies were resolved at 2.59 Å, 2.59 Å, and 2.54 Å resolution, respectively **(Fig. 2b-d and Extended Data Fig. 3 and 4)**. Both the dimeric and tetrameric assemblies exhibit C2 symmetry, whereas the trimeric population displays an asymmetric architecture **(Fig. 2b)**.

Structural comparison of the dimeric, trimeric, and tetrameric assemblies reveals a conserved butterfly-shaped core dimer that is maintained across all oligomeric states **(Fig. 2b)**. This core represents the conserved structural unit that persists unchanged upon higher-order assembly. The trimeric population does not form a symmetric trimer; instead, it corresponds to a partially assembled intermediate in which an additional protomer associates asymmetrically with one side of the dimeric assembly. Completion of this association on both sides yields the symmetric tetrameric assembly. In sum, these observations suggest a stepwise, modular plug-in oligomerization, in which a stable core dimer serves as a structural scaffold for the sequential recruitment of functional protomeric units, while the observed oligomeric heterogeneity could be concentration dependent.

Despite the different oligomeric states, the overall architecture remained highly conserved, with the individual HoxEFUYH protomers and the core dimer preserved across the dimeric, trimeric, and tetrameric assemblies **(Fig. 2b-d)**. In contrast to the apo HoxEFUYH_2_ reconstruction, all holo structures displayed well-defined density for both FMN and NADH within the reductase module, indicating that cofactor binding does not induce major conformational rearrangements. Because the tetrameric reconstruction represents the highest-order assembly while retaining the same protomer organization and cofactor arrangement, all subsequent structural analyses are based on the tetrameric HoxEFUYH_4_ structure unless stated otherwise **(Fig. 2c,d)**.

### Electron transfer pathway and [NiFe] active site

To define the molecular basis of electron transfer in HoxEFUYH, we next analyzed the cofactor architecture of the holo enzyme. Across each protomer, these cofactors define a continuous electron-transfer pathway linking the reductase and hydrogenase units **(Fig. 2b-d)**. Consistent with this structural organization, consecutive FeS clusters are positioned with edge-to-edge distances below 14 Å, supporting rapid electron tunneling^36,37^ **(Fig. 3a)**. The [NiFe] active site is electronically connected to the proximal Y1 cluster, which in turn links to the chain of FeS cluster within HoxU and the reductase module, establishing a continuous electron transfer pathway across the entire protomer. The reductase module is organized around a conserved HoxEFU core that is structurally related to the HydBC modules of electron-bifurcating hydrogenases and transhydrogenases, as well as to the N-module of respiratory complex I^23–26,38,39^. The HoxE subunit is homologous to the HydC subunit of HydBC and to Nqo2 of complex I **(Extended Data Fig. 5a,c)** and comprises of an N-terminal domain followed by a C-terminal thioredoxin-like domain that coordinates a [2Fe-2S] cluster (E1) via four conserved cysteine residues **(Fig. 3a and Extended Data Fig. 5a,b)**. On the other hand, HoxF shows close structural similarity to the HydB subunit of HydBC and to Nqo1 of complex I **(Extended Data Fig. 5a,c)**. However, unlike HydB, HoxF lacks the C-terminal ferredoxin-like domain that harbors additional FeS clusters in bifurcating hydrogenases, indicating a distinct organization of the electron-transfer network in HoxEFUYH. Its N-terminal region harbors a [2Fe-2S] cluster (F2) and is followed by a Rossmann-fold region and an adjacent ubiquitin-like domain that together form the binding site for FMN and NADH. The C-terminal region of HoxF coordinates an [4Fe-4S] cluster (F1), which is positioned at the interface to HoxU **(Fig. 3a and Extended Data Fig. 5a-c)**.

**Fig. 3:**
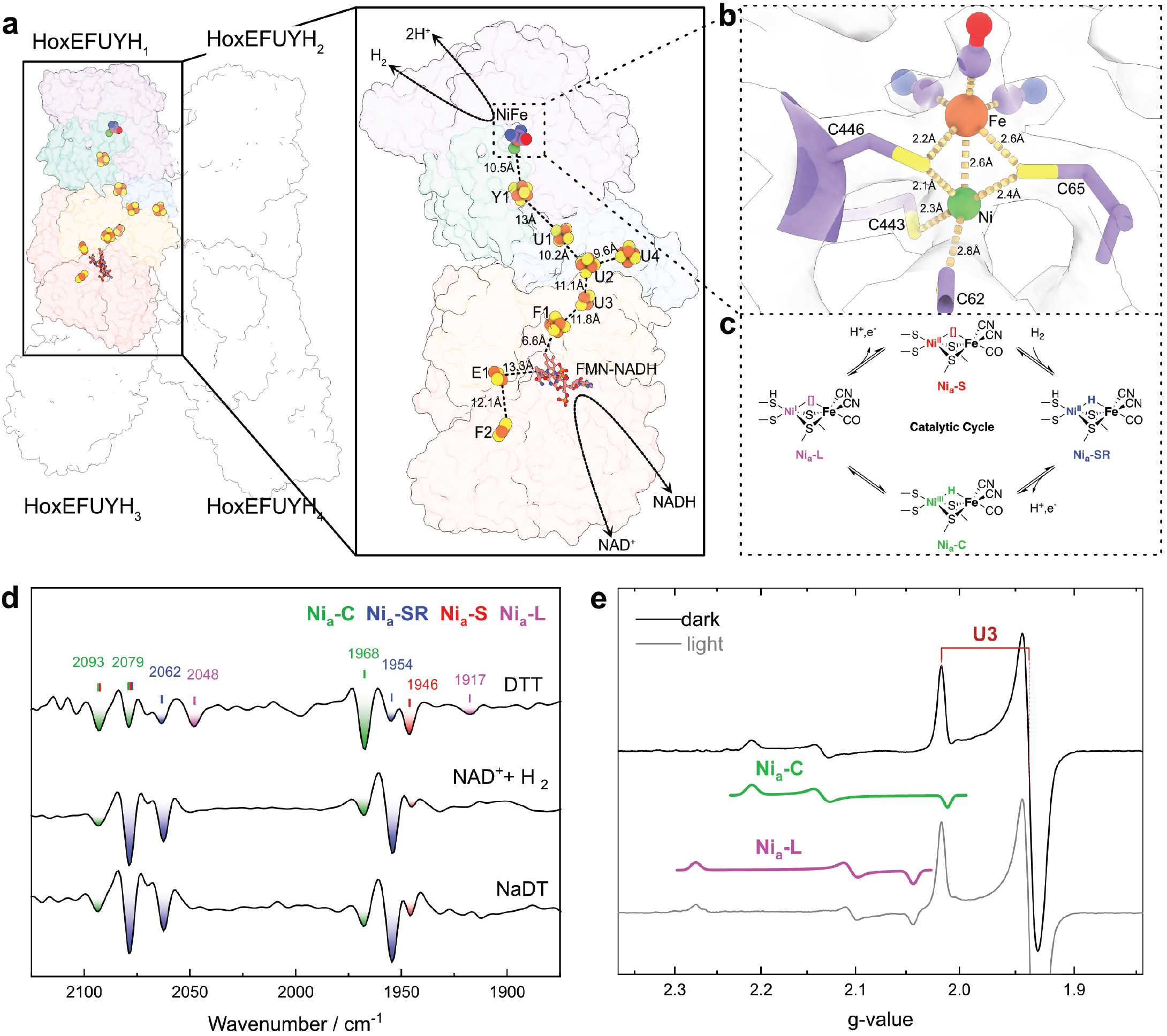
Cofactor arrangement, [NiFe] active site architecture, and catalytic intermediates of the HoxEFUYH hydrogenase. **a**, Arrangement of the redox cofactors and electron-transfer pathway within a single HoxEFUYH protomer. The enlarged view highlights the [NiFe] active site, FMN-NADH binding site, and the FeS relay connecting the hydrogenase and reductase modules. Edge-to-edge distances between adjacent cofactors are indicated. **b**, Close-up view of the catalytic [NiFe] active site coordinated by conserved cysteine residues within HoxH. Distances between the metal ions and coordinating ligands are indicated. **c**, Proposed catalytic cycle of the [NiFe] active site showing the major catalytic intermediates identified spectroscopically, including Ni_a_-S, Ni_a_-SR, Ni_a_-C, and Ni_a_-L. **d**, second derivatives of IR spectra from HoxEFUYH taken under different redox conditions exhibiting characteristic CO and CN stretching frequencies, which correspond to distinct catalytic intermediates of the [NiFe] active site. Spectroscopic data obtained under DTT-reduced, NAD^+^ + H_2_, and NaDT-reduced conditions are displayed here. **e**, EPR spectra of DTT-reduced Hox-EFUYH recorded at 35 K showing the Fe-S cluster signal assigned to U3 and light-driven conversion between the Ni_a_-C and Ni_a_-L states of the [NiFe] active site.

**Fig. 4:**
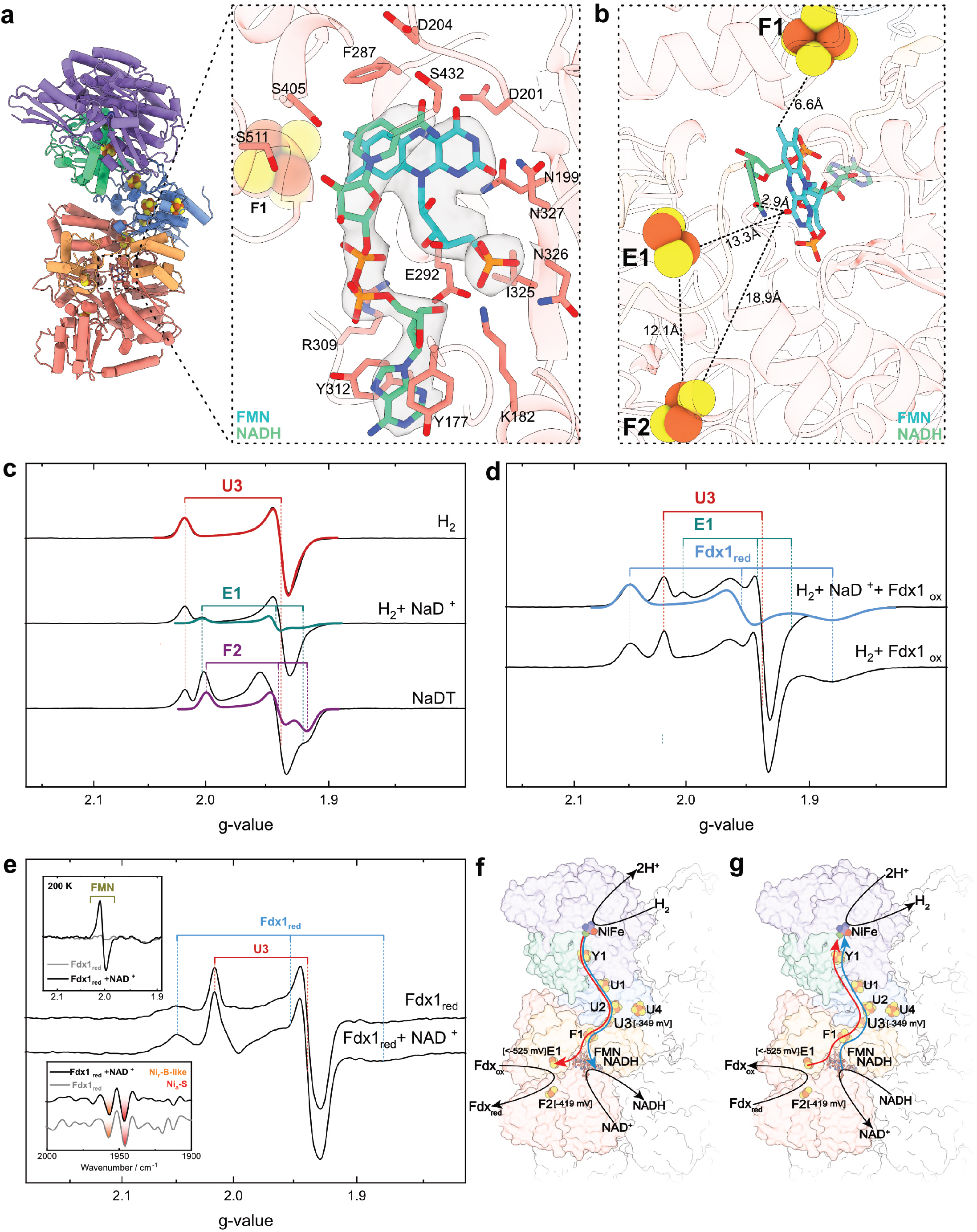
FMN-NADH binding site, FeS cluster spectroscopy, and proposed electron-transfer pathway in HoxEFUYH. **a**, Cryo-EM density of the FMN-NADH binding site within HoxF. The enlarged view highlights the coordination environment surrounding FMN and NADH. **b**, FMN-NADH binding site and its connectivity to the neighboring FeS clusters F1, E1, and F2. Edge-to-edge distances between adjacent cofactors are indicated. **c**, EPR spectra of HoxEFUYH under different reducing conditions showing signals assigned to the U3, E1, and F2 FeS clusters. Spectra obtained under H_2_, H_2_ + NAD^+^, and NaDT-reduced conditions are shown. **d**, EPR spectra recorded in the presence of oxidized ferredoxin (Fdx1_ox_) showing reduction of Fdx1 together with signals from the U3 and E1 FeS clusters during H_2_ oxidation. **e**, EPR spectra of reduced ferredoxin (Fdx1_red_) in the absence and presence of NAD^+^. Insets show the characteristic FMN radical signal and corresponding second derivatives of IR spectra from the [NiFe] active site under the indicated redox conditions. All EPR spectra shown were recorded at 35 K, if not indicated otherwise. **f**,**g**, Proposed electron-transfer pathways within HoxEFUYH during electron bifurcation and confurcation. During H_2_ oxidation **(f)**, electrons derived from H_2_ at the catalytic [NiFe] active site are split into a high-potential branch that drives NAD^+^ reduction and a low-potential branch that drives ferredoxin reduction. During H_2_ evolution **(g)**, electrons supplied by NADH and reduced ferredoxin enter the high-potential and low-potential branches, respectively, and converge at the FMN-containing reductase module before being transferred through the FeS relay to the [NiFe] active site, where proton reduction generates H_2_. Colored pathways indicate electron flow through the high-potential (blue) and low-potential (red) branches.

**Fig. 5:**
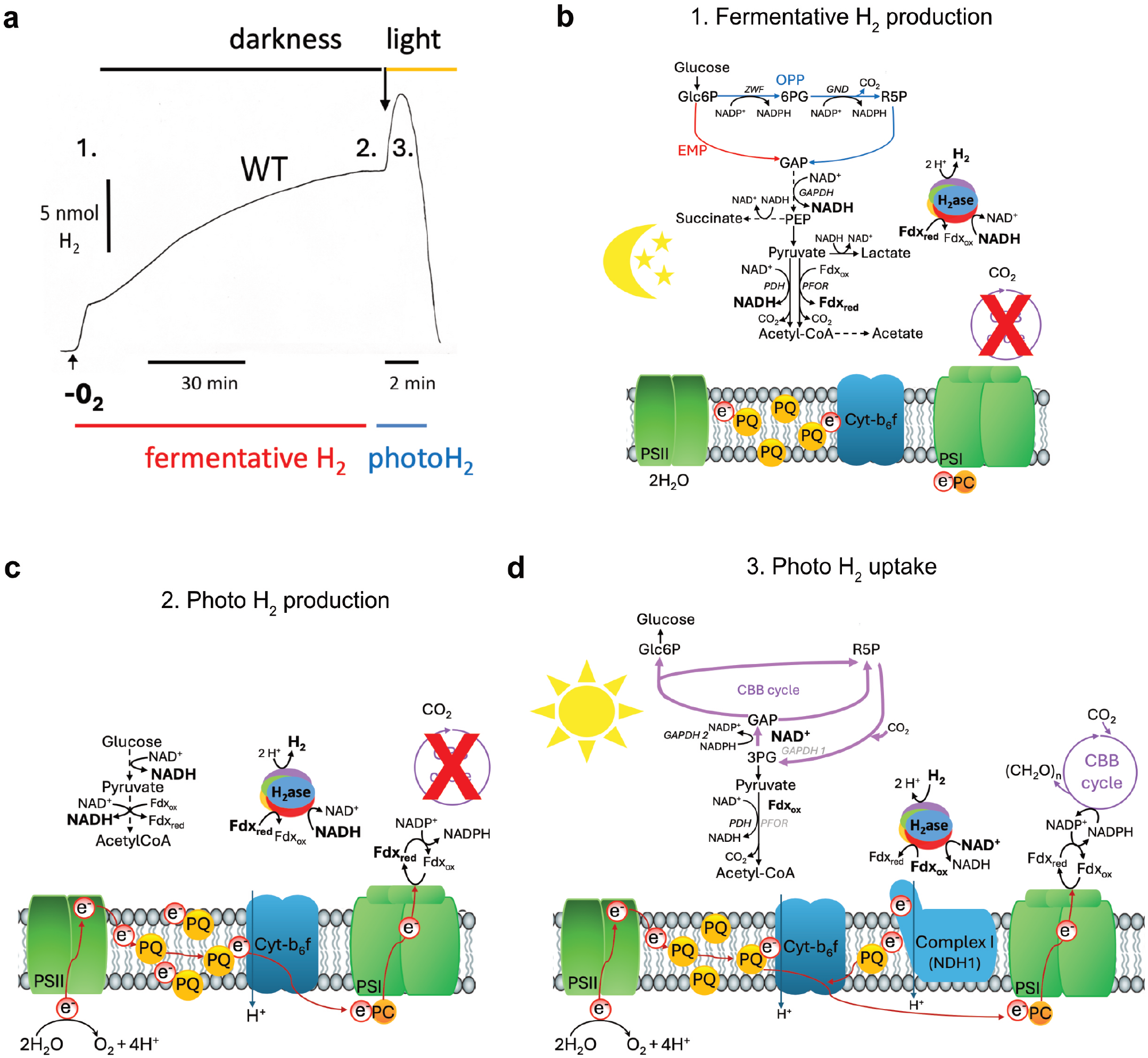
Proposed physiological role of HoxEFUYH in cyanobacterial H2 metabolism. **a**, Representative H_2_ production and uptake profile measured during the transition from dark anaerobic conditions to illumination. Under darkness, cells accumulate fermentative H_2_, whereas illumination initially stimulates photoH_2_ production followed by rapid H_2_ uptake. **b**, Metabolic pathways supporting fermentative H_2_ production under dark anaerobic conditions. Carbohydrates are oxidized through the oxidative pentose phosphate pathway (OPP), Embden-Meyerhof-Parnas pathway (EMP), lower glycolysis, pyruvate dehydrogenase (PDH), and pyruvate:ferredoxin oxidoreductase (PFOR), generating high levels of NADH and reduced ferredoxin (Fdx_red_). Production of lactate, succinate, acetate, and H_2_ regenerates NAD^+^ and oxidized ferredoxin (Fdx_ox_). **c**, Metabolic state during photoH_2_ production immediately after the onset of illumination. NADH levels remain high from the preceding dark period, while Fdx_red_ accumulates at PSI because the Calvin-Benson-Bassham (CBB) cycle is not yet fully active. The simultaneous availability of NADH and Fdx_red_ promotes H_2_ production by Hox-EFUYH. **d**, Metabolic state during photoH_2_ uptake under prolonged illumination. Activation of the CBB cycle increases gluconeogenetic carbon flux and consumption of reducing equivalents. GAPDH2 becomes active and operates in the opposite direction of GAPDH1, promoting oxidation of NADPH, while NAD^+^ is no longer reduced by GAPDH1, leading to decreased NADH levels in the light. PFOR is likely inactive under these conditions. Together with elevated H_2_ concentrations, these metabolic changes favor H_2_ uptake by HoxEFUYH.

In contrast, the associated subunit HoxU is structurally distinct and does not represent a direct homologue of HydA, but it shows similarities with the HoxU subunit from the non-bifurcating HoxFUYH hydrogenase (RMSD 0.9Å) from *Hydrogenophilus thermoluteolus*^*40*^ with RMSD value of 0.9Å **(Extended Data Fig. 5e)**. HoxU comprises an N-terminal [FeFe]-hydrogenase–like fold and a C-terminal extension that together coordinate three [4Fe-4S] clusters (U1, U2 and U4), as well as a single [2Fe-2S] cluster (U3) **(Fig. 3a and Extended Data Fig. 5a,b)**. The clusters U1, U2 and U4 and the overall fold is also very similar compared to the Nqo3 subunit of the complex I of *Thermus thermopilus*^*39*^. Notably, the U1 cluster is coordinated by a ligand sphere having three cysteines and one histidine, distinguishing it from the canonical all-cysteine coordination observed for the other FeS clusters **(Extended Data Fig. 5b)**. This positions HoxU as the conduit between the reductase and hydrogenase modules. Together, this arrangement establishes a continuous chain of FeS cluster network connecting the FMN/NADH-binding site in HoxF with the hydrogenase module via HoxU, providing the structural basis for long-range ET **(Fig. 3a)**^41,42^.

Extending this organization into the hydrogenase module, the [NiFe] active site in HoxH is positioned in close proximity to the proximal Y1 [4Fe-4S] cluster in HoxY **(Fig. 3a)**. HoxY shows high structural similarity to the HydS subunit of the bifurcating [NiFe]-hydrogenase HydABCSL complex from *Acetomicrobium mobile*^*25*^ and HoxY subunit from the non-bifurcating HoxFUYH hydrogenase from *H. thermoluteolus*^40^ **(Extended Data Fig. 5e,f)**. Like its homologs, HoxY coordinates a [4Fe-4S] cluster (Y1) **(Fig. 3a and Extended Data Fig. 5a,b)**. Whereas HoxH closely resembles the HydL subunit of HydABCSL **(Extended Data Fig. 5f)** and harbors the catalytic [NiFe] active site. The spatial arrangement of the [NiFe] cofactor relative to the proximal FeS cluster establishes a direct connection between the catalytic center and the extended FeS network. Notably, the Ni-Fe distance is ~2.6 Å, consistent with a canonical [NiFe]-hydrogenase active site geometry **(Fig. 3b)**. Furthermore, at the local resolution of ~2.3 Å, no density corresponding to a bridging oxygen (e.g., hydroxide) ligand is observed between the two metals, in agreement with Fourier-transform infrared (FTIR) spectroscopic data **(Fig. 3d)**.

To assess whether the structurally defined active site is catalytically engaged, we monitored the redox states of the Ni-Fe center by FTIR and electron paramagnetic resonance (EPR) spectroscopy. Under the same mildly reducing conditions used for the cryo-EM experiments (DTT, E_0_,SHE ≈ –330 mV), second derivative IR spectra reveal a mixture of all four catalytic intermediates (Ni_a_-S, Ni_a_-SR, Ni_a_-C, and Ni_a_-L), assigned by their characteristic CO and CN stretching frequencies^19^ **(Fig. 3c, d; Supplementary Table 3)**. Using the physiological substrates H_2_ and NAD^+^ or the chemical reductant sodium dithionite (NaDT), the active site could be further reduced, enriching the Ni_a_-SR state, which forms a Ni_r_-B-like intermediate, when being exposed to O_2_ **(Extended Data Fig. 10a)**. Ni_a_-C, the predominant species under DTT, was detected by EPR at 80 K for the first time in this enzyme and could be reversibly converted into its tautomer Ni_a_-L upon illumination **(Fig. 3e)**. Notably, the spectral intensity of these two species accounts for less than 10% of the total [NiFe] active site population **(Extended Data Fig. 9a)** in the EPR spectrum, whereas these species contribute more than 50% of the IR intensity **(Fig. 3d)**. Because the spectra were recorded at 283 K (IR) and 80 K (EPR), the most plausible explanation is a temperature-dependent shift in the redox equilibrium toward enrichment of the fully reduced Ni_a_-SR state at lower temperature. A similar behavior has been reported for the structurally related [NiFe]-hydrogenase from *Hydrogenophilus thermoluteolus*^43^. This observation indicates thermally driven electron redistribution among the cofactors without substantial energy barriers, consistent with the bidirectional function of the enzyme.

### Structural and spectroscopic basis of electron bifurcation at the FMN site

To examine how electrons enter and are distributed within the complex, we next studied the FMN/NADH-binding site and the associated FeS network in the structure **(Fig. 4a)**. Binding of FMN and NADH does not induce any large-scale conformational changes in the HoxEFUYH_4_ holo state, indicating that the reductase module is pre-organized for substrate (or coenzyme) engagement. Instead, FMN/NADH binding is accommodated by localized side-chain rearrangements within the active site. Comparison of the apo and holo structures reveals a pronounced reorientation of Phe287 upon FMN/NADH binding, accompanied by smaller adjustments of Tyr177 and Tyr312 to create a binding pocket for the adenine moiety of NADH **(Extended Data Fig. 6a)**.

The microenvironment of the FMN/NADH site is defined by a network of conserved residues within 4 Å of the substrates **(Fig. 4a; Extended Data Fig. 6a-c)**. FMN is primarily stabilized through electrostatic interactions involving Lys182, Asn199, Asn326, and Asn327, whereas Tyr177 and Tyr312 form a π-stacking arrangement around the adenine ring of NADH **(Fig. 4a)**. Notably, Asp201 is positioned near the isoalloxazine ring of the FMN and may modulate the redox properties of the flavin through local electrostatic interactions. Consistent with its functional role, the FMN/NADH-binding region of HoxF contains the three conserved signature motifs characteristic of bifurcating hydrogenases: the Ala-Phe-Met (AFM) motif, the single-phenylalanine motif, and the Gly-Gly-Pro-Ser-Gly (GGPSG) motif, all of which contribute to shaping the flavin-binding pocket. All three motifs are preserved in Hox and occupy similar positions to those observed in the holo structures of HydABC and HydABCSL^23,25^ **(Extended Data Fig. 6c)**. Comparison of the apo and holo structures reveals loops spanning residues 201-207 and 307-322 exhibit localized rearrangements upon NADH binding **(Extended Data Fig. 6b)**. Together these similarities underscore the structural and functional conservation between the HoxEFU reductase module and the HydBC-type bifurcation machinery.

We next analyzed the spatial arrangement of the FMN/NAD(H) site and the surrounding FeS clusters to understand how electrons are transferred through the reductase module **(Fig. 4b)**. The FMN is positioned between the NADH-binding site and the neighboring F1 and E1 FeS clusters, forming the immediate redox environment of the flavin. The distance between the NADH nicotinamide and the FMN isoalloxazine ring is approximately 2.9 respectively)^46^. These findings are corroborated by experiments with the strong chemical reductant NaDT (E_0,SHE_ = −660 mV)^47^. Under these conditions, the [4Fe-4S] cluster signals increase only moderately, whereas the F2 and E1 clusters become fully reduced **(Figure 4c and Extended Data Fig. 9b)**, consistent with their substantially lower reduction potentials.

To directly probe electron transfer into the low-potential branch, we next examined the interaction of HoxEFUYH with ferredoxin. The oxidized enzyme was incubated with Å, allowing direct hydride transfer and generation of reduced flavin states. The reduced flavin can subsequently distribute electrons via sequential single-electron transfer reactions to the surrounding FeS network^44,45^. Both F1 and E1 lie within electron-transfer distance of FMN, making them the primary acceptors of electrons from the flavin. In contrast, the much larger distance between FMN and the F2 [2Fe-2S] cluster (~19 Å) precludes direct electron transfer, thereby enforcing electron flow through the proximal F1 and E1 clusters rather than direct exchange with F2 **(Fig. 4b)**. The terminal [2Fe-2S] clusters E1 and F2 are positioned close to the protein surface and are surrounded by several solvent-exposed Lys and Arg residues that generate locally electropositive surface patches. These features are consistent with a potential interaction interface for negatively charged ferredoxin molecules and position E1 and F2 as likely sites of electron exchange between HoxEFUYH and soluble electron carriers **(Extended Data Fig. 6d**,**e)**.

To gain insight into the electronic landscape of the cofactors during electron bifurcation and confurcation, we incubated oxidized HoxEFUYH with different combinations of its physiological substrates and recorded EPR spectra. Upon exposure to H_2_, the characteristic spectrum of the [2Fe-2S] cluster U3 (E_m,SHE_ = −349 mV)^38^ and signals from at least four different [4Fe-4S] clusters can be observed^6,46^ **(Fig. 4c and Extended Data Fig. 7c)**. Addition of NAD^+^ results in the appearance of an additional [2Fe-2S] cluster signal, assigned to E1 based on its spectral features **(Extended Data Fig. 9c and 10c)**. Notably, this indicates that the binding of NAD^+^ at the FMN enables the electrons generated by H_2_ oxidation to be transferred also to E1, supporting the assignment of the flavin as the site for bifurcation.

However, under H_2_ and NAD^+^, electrons still appear to accumulate preferentially within the FeS relay of the high-potential NAD(H)-linked branch, whereas the F2 and E1 clusters of the low-potential ferredoxin branch remain largely oxidized. This is consistent with their lower mid-point potentials verus SHE (−419 mV and < −525 mV, oxidized ferredoxin 1 (Fdx1_ox_) in the presence of H_2_ and NAD^+^ **(Fig. 4d)**. The EPR spectrum displays an additional signal attributable to reduced Fdx1, confirming H_2_-driven ferredoxin reduction in agreement with biochemical assays **(Fig. 1e)** and consistent with electron transfer via the low-potential branch. The EPR signal and power saturation curve **(Extended Data Fig. 8f and 10d)** of this signal is essentially identical to the one of the isolated Fdx1 and shows no magnetic coupling to other paramagnetic centers, which supports a transient interaction of Fdx1 with HoxEFUYH rather than a stable binding. Consistent with this, biochemical assays showed NAD^+^ reduction during H_2_ oxidation in the presence of Fdx1. Notably, a signal corresponding to reduced Fdx1 is also observed in the absence of NAD^+^ **(Fig. 4d)**, suggesting partial electron leakage or reduction by dissociated HoxH-containing species that retain the catalytic [NiFe] active site.

To assess the confurcation pathway, we next tested whether reduced ferredoxin can inject electrons into the HoxEFUYH electron-transfer network. When oxidized HoxEFUYH was exposed to a threefold excess of Fdx1_red_ **(Fig. 4e and Extended Data Fig. 10e)**, the EPR signal of U3 and the IR signature of the most oxidized catalytic Ni_a_-S state are observed, indicating that electrons can enter the enzyme via the ferredoxin-linked branch but do not substantially reduce the [NiFe] active site **(Extended Data Fig. 10a)**. These observations demonstrate electronic communication between ferredoxin and the internal FeS relay. However, consistent with the biochemical assay **(Fig. 1b)**, electron input from Fdx1_red_ alone is insufficient to drive efficient H_2_ production, indicating that additional reducing equivalents from NADH are required for confurcation. Strikingly, addition of NAD^+^ to the reaction mixture enabled detection of a flavin semiquinone radical **(Fig. 4e)**, which appears to be stabilized only under these electron-limiting conditions. This observation further supports a central role of FMN as the electron-bifurcation/confurcation center and indicates that the flavin semiquinone is preferentially stabilized in the presence of NAD^+^.

Together, the structural and spectroscopic data support a model in which the FMN site as the central hub connecting electronically distinct high- and low-potential branches within HoxEFUYH. During bifurcation, electrons derived from H_2_ oxidation are transferred from the catalytic [NiFe] site through the FeS relay to the FMN, where they are bifurcated toward NAD^+^ reduction (high-potential) and the low-potential E1/F2 branch involved in ferredoxin reduction **(Fig. 4f)**. Conversely, during confurcation, electrons originating from NADH and Fdx_red_ converge at the FMN site and are transferred through the FeS relay toward the catalytic [NiFe] center to drive H_2_ production **(Fig. 4g)**.

## Discussion

In this study, we have achieved the first isolation and comprehensive characterization of the intact, bidirectional [NiFe]-hydrogenase HoxEFUYH from *Synechocystis* sp. PCC 6803, defining its structure, redox properties, and catalytic behavior. We demonstrate that the enzyme forms a modular oligomeric assembly capable of both bifurcating and confurcating electron flow between H_2_, NAD(H), and ferredoxin, thereby reconciling previously conflicting biochemical data on donor specificity. Together, our data reveal a connected ET network that is structurally organized for bidirectional catalysis. Our high-resolution cryo-EM structures show a continuous ET pathway linking the FMN/NADH-binding site in the reductase module to the [NiFe] catalytic center in the hydrogenase module via a series of closely spaced FeS clusters. Spectroscopic investigations under physiologically relevant and controlled redox conditions corroborate this pathway and support its role in bidirectional catalysis.

Mechanistically, our data demonstrate that the cyanobacterial [NiFe]-hydrogenase HoxEFUYH organizes electron flow in a manner that is reminiscent of anaerobic electron-bifurcating HydABC complexes, nevertheless differs substantially in architecture and regulatory strategy. In canonical HydABC systems from obligate anaerobes such as *Acetobacterium woodii* and *Thermoanaerobacter kivui*, bifurcation is achieved through redox-controlled conformational gating. Reduction of the proximal B2 [2Fe-2S] cluster, which is homologous to the F2 cluster in HoxEFUYH, increases the affinity for NAD(P)^+^, thereby enabling exergonic hydride transfer at the flavin site. Subsequent nucleotide reduction and release trigger repositioning of the mobile C-terminal HydC domain, allowing its [2Fe-2S] cluster to engage the low-potential ferredoxin branch. These redox-driven structural rearrangements establish a kinetic barrier that prevents electron backflow from reduced ferredoxin to the FMN site. Thus, energy coupling in HydABC emerges from conformational dynamics imposed by redox chemistry rather than from intrinsic flavin thermodynamics.

In contrast, HoxEFUYH lacks the defining architectural features required for such a gating mechanism. The complex does not contain a tethered C-terminal ferredoxin domain comparable to HydC, nor does it harbor a permanently associated bacterial-type ferredoxin. Instead, the cyanobacterial enzyme interfaces with soluble, plant-type [2Fe-2S] ferredoxins, reflecting its integration into photosynthetic and fermentative redox metabolism^34,48,49^. Consequently, the classical embedded ferredoxin module observed in anaerobic bifurcating systems is absent, implying that electron donation must occur via an alternative, externally accessible binding surface.

An outstanding question therefore concerns the location and nature of the ferredoxin binding site. Functional re-purposing within the cyanobacterial complex may involve the HoxE subunit, although no canonical docking pocket is structurally apparent. Comparison with bifurcating oxidoreductases suggests a potential interaction interface near the N-terminal region of the HydB-like subunit HoxF^38^, while the electropositive surface surrounding E1/F2 may facilitate the transient electrostatic interactions required for rapid and reversible docking. Consistent with this model, attempts to capture a stable HoxEFUYH-ferredoxin complex were unsuccessful. Moreover, reduced ferredoxin detected under turnover conditions shows no magnetic coupling to internal FeS clusters, further supporting a transient interaction mechanism. Unlike canonical HydABC hydrogenases, HoxEFUYH therefore appears not to rely on redox-dependent movements of tethered ferredoxin-like domains.

These mechanistic insights also provide a framework for understanding the physiological role of HoxEFUYH during fermentative and photosynthetic metabolism. In the dark, carbohydrates are primarily oxidized via the oxidative pentose phosphate pathway, producing NADPH^50,51^. NADH, on the other hand, is formed in lower-glycolysis via glyceraldehyde-3-phosphate dehydrogenase (GapDH1, slr0884). GapDH2 (sll1342), which can utilize both NAD(H) and NADP(H), is primarily involved in the CBB cycle and is inactivated in the dark^20^. The pyruvate dehydrogenase complex (PDH) of *Synechocystis*, which supplies NADH, is inactivated at high NADH concentrations and is functionally replaced by pyruvate:ferredoxin oxidoreductase (PFOR), which produces Fdx_red_^34^. Thus, dark fermentative metabolism inherently generates NADH and Fdx_red_ through distinct but connected pathways.

Initially it was assumed that fermentative H_2_ production relies on NAD(P)H due to the homology of HoxF to the NADH binding module of photosynthetic complex I (NDH-1). *In vitro* assays supported this assumption. In line with this, deletion of NADH consuming enzymes as e.g. lactate dehydrogenase resulted in enhanced fermentative H_2_ production^52^. These observations were later questioned by thermodynamic calculations indicating that measured H_2_ concentrations could not be satisfactorily explained by physiological NAD(P)H levels^16^. This, in turn, shifted attention to ferredoxin as a possible reductant^16^.

Subsequent *in vitro* assays using cell homogenates, dithionite-reduced ferredoxin, and Clark-type electrodes, together with the reduced fermentative H_2_ production observed in a PFOR knockout strain (Δ*pfor*), suggested Fdx_red_ as electron donor for H_2_ production^16^. However, this interpretation was later challenged when Clark-type electrodes were found to produce signals even in cell homogenates of a hydrogenase-free (Δ*hox*) strain in the presence of dithionite and ferredoxin, indicating potential measurement artifacts^17^. The confurcating nature of HoxEFUYH resolves these previously conflicting observations: efficient H_2_ production requires both NADH and Fdx_red_, thereby reconciling earlier biochemical data with thermodynamic considerations. The redox partners NADH and Fdx_red_ therefore connect fermentative H_2_ production to lower glycolysis and pyruvate decarboxylation **(Fig. 5a, b)**.

Oxidative decarboxylation of pyruvate generates acetyl-CoA and allows the synthesis of additional ATP per acetate. Under dark, anaerobic glucose degradation, cells can generate up to four ATP per glucose via acetate and CO_2_ formation, effectively doubling the energy yield relative to lactic acid or ethanol fermentation. The concomitant formation of two NADH molecules and four reduced ferredoxins via PFOR per glucose is matched precisely by the confurcating activity of the bidirectional [NiFe]-hydrogenase, which can convert these reductants into H_2_ **(Fig. 5a, b)**. Thus, the enzyme operates not only as a provider of highly reducing equivalents but also as an energy-conserving element under fermentative conditions.

The specificity for NADH over NADPH in combination with reduced ferredoxin provides further physiological context. During the transition from dark anaerobiosis to light, both NADH and reduced ferredoxin are initially present, allowing H_2_ production to rise briefly upon illumination **(Fig. 5a, c)**. However, because the NADH pool is limited and is not efficiently replenished by photosynthetic metabolism, H_2_ evolution is expected to cease rapidly once this pool is consumed, thereby preventing sustained and wasteful energy loss. Conversely, H_2_ uptake via the soluble redox partners alone appears less likely under early light conditions, because PSI should rapidly reduce ferredoxin, while NADH becomes progressively oxidized. *In vivo*, an alternative route for H_2_ uptake has therefore been proposed in which HoxEFUYH may interact with the cyanobacterial photosynthetic complex I, which lacks NADH-binding subunits, and transfers electrons toward plastoquinone. In this case NAD^+^ and Fdx_ox_ might not be required *in vivo*. Structural features, such as the additional U4 [4Fe-4S] cluster and the arrangement of Fe-S clusters in the diaphorase module, support this hypothesis by analogy with complex I. However, the existence and physiological relevance of such a Hox-NDH-1 connection remains unresolved and should presently be regarded as speculative. Along this line, it has been shown that both the NDH complex and the photosynthetic electron transfer between the PQ pool, the CytB_6_f complex, and PSI are necessary for proper photoH_2_ uptake^17,21^.

Based on our results, photoH_2_ production in wild-type strains of *Synechocystis* must be reevaluated. Until now, it was widely believed that photoH_2_ production, whether via Fdx_red_ or NADPH from PSI, relies exclusively on the photosynthetic electron transport chain^16,17^. However, our data show that the electrons used in photoH_2_ production never originate exclusively from the light reaction; rather, half of them come from carbohydrate oxidation in the form of NADH. From a biotechnological perspective, photoH_2_ production in wild-type cyanobacteria is therefore less attractive than previously assumed. This illustrates and underscores that the approach of directly coupling the hydrogenase HoxYH to PSI without the diaphorase subunit (HoxEFU) is the most direct way to utilize electrons from photosynthesis exclusively for the production of photoH_2_^2,27^.

Building on all these insights, the structural and mechanistic understanding of [NiFe]-hydrogenase HoxEFUYH opens new avenues for biotechnological innovation in green H_2_ production. The transient, solvent-exposed ferredoxin interface in HoxEFUYH, together with its modular diaphorase architecture, provides a structural blueprint for engineering more efficient ET pathways or optimized docking with PSI.

## Methods

### Mutant strain generation and *in vivo* hydrogenase activity

All amplificants were produced with primers listed in table S1 and assembled via TAR cloning^53^. For homologous recombination in *Synechocystis*, a plasmid backbone with recombination sites for the *hox* locus (pMQ80hox) was generated first and subsequently restricted with SmiI (SwaI) to construct pP_psbAII_*Syn*Hox-TST. Plasmid integrity was verified via Sanger sequencing (GENEWIZ, Germany GmbH, Leipzig). A *Synechocystis* strain lacking all *hox* genes (*Δhox*) was then transformed with pP_psbAII_*Syn*Hox-TST. To screen clones for hydrogenase activity, methyl viologen assays (5 mM methyl viologen, 20 mM sodium dithionite) were performed via gas chromatography (Nexis^™^ GC-2030, Shimadzu, Kyoto, Japan). One clone with similar activity levels to the WT was selected from this assay to monitor *in vivo* fermentative and photosynthetic H_2_ production under anaerobic conditions (10 mM glucose, 4 U/µl glucose oxidase, 5 U/µl catalase) using Clark-type H_2_ sensors (Unisense, Aarhus, Denmark), which also showed similar activity to the WT **(Fig. 1 and Extended Data Fig. 1b,c)**.

### *In vitro* hydrogenase activity (biochemistry)

Hydrogenase activity of isolated hydrogenase was measured under anaerobic conditions (40 mM glucose, 16 U/µl glucose oxidase, 20 U/µl catalase) using a Clark-type H_2_ sensor (Unisense, Aarhus, Denmark) combined with scanning absorption between 340 and 460 nm (Cary 60 UV-Vis Spectrophotometer) for simultaneous monitoring of NAD(P)H and ferredoxin redox changes. For H_2_ production experiments, pyruvate-ferredoxin-oxidoreductase (PFOR) was used for reduction of ferredoxin. 2.5 µM PFOR were activated prior to addition by incubation for at least 40 minutes at RT with substrate and cofactors under anaerobic conditions (100 µM Tris pH 8, 5 mM TPP, 1 mM CoA, 10 mM pyruvate, 40 mM glucose, 16 U/µl glucose oxidase, 20 U/µl catalase). In a 2 ml cuvette, 0.1-0.8 µM hydrogenase were added to a buffer mix (100 mM Tris pH 8, 5 mM TPP, 1 mM CoA, 10 mM pyruvate, 40 mM glucose, 16 U/µl glucose oxidase, 20 U/µl catalase) containing 0.125 µM active PFOR. To induce H_2_ production, 10 µM ferredoxin and up to 100 µM NAD(P)H were added. For H_2_ uptake experiments, the buffer mix (100 mM Tris pH 8, 40 mM glucose, 16 U/µl glucose oxidase, 20 U/µl catalase) was prepared with 25% H_2_-saturated water (corresponding to 191 µM H_2_ at 28°C). 10-20 µM oxidized ferredoxin and 1 mM NAD(P)^+^ were added as electron acceptors to induce H_2_ uptake.

### Expression and purification of HoxEFUYH hydrogenase from *Synechocystis* PCC 6803

Cells of *Synechocystis* sp. PCC 6803 expressing Twin-Strep-tagged HoxH were cultivated under standard photo-autotrophic conditions and harvested by centrifugation at 8000×g for 10 min at 4°C. All subsequent purification steps were carried out under strictly anaerobic conditions (95-98% N_2_ and 2-5% H_2_) to preserve enzyme activity. Cell pellets were resuspended in lysis buffer (100 mM Tris-HCl pH 8.0, 150 mM NaCl, 1 mM EDTA, 2 mM DTT) supplemented with DNase I and protease inhibitors. Cells were lysed using two passes through a French press at 20,000 psi. Cell debris and unbroken cells were removed by centrifugation at 25,000 rpm for 1 h using a Ti-25.15 rotor. The soluble fraction (supernatant) containing the HoxEFUYH complex was loaded onto a Strep-Tactin affinity column pre-equilibrated with lysis buffer. After extensive washing to remove non-specifically bound proteins, the hydrogenase complex was eluted with 2.5 mM desthiobiotin. Eluted fractions were concentrated and further purified by size-exclusion chromatography (SEC) using a Superose 6 Increase column equilibrated in SEC buffer (100 mM Tris-HCl pH 8.0, 150 mM NaCl, 1 mM EDTA, 2 mM DTT). Peak fractions corresponding to the intact HoxEFUYH complex were pooled and analyzed by SDS-PAGE to assess purity. Protein concentration was determined using Bradford assay, and samples were always kept under anaerobic conditions. The purified enzyme was either used immediately for biochemical assays, cryo-EM grids preparation or flash-frozen in liquid nitrogen and stored at −80°C until further use.

### Cryo-EM sample preparation, data acquisition, processing and model building

For cryo-EM grid preparation, purified HoxEFUYH hydrogenase was handled under strictly anaerobic conditions (95-98% N_2_ and 2-5% H_2_). For the apo sample, freshly purified protein was directly used, whereas for the holo sample, the enzyme was incubated with NADH (500 µM) and FMN (50 µM) prior to vitrification. In addition, 1 mM of CHAPSO was included for the holo condition to improve particle distribution and reduce preferred orientation. 4 µl of protein was applied to glow-discharged Quantifoil R 2/1 copper 300-mesh grids (PELCO easiGlow, 25 s at 15 mA). Grids were blotted and plunge-frozen in ethane-propane using a Vitrobot Mark IV (Thermo Fisher Scientific) at 4°C and 100% humidity under anaerobic conditions. Cryo-EM data were collected using Smart EPU on a Titan Krios G4 transmission electron microscope (Thermo Fisher Scientific) operated at an accelerating voltage of 300 kV. The microscope was equipped with a Selectris energy filter using a 10-eV slit width and a Falcon 4i direct electron detector. Data were recorded in counting mode at a nominal magnification of 165,000×, corresponding to a calibrated pixel size of 0.73 Å. Images were acquired with a total electron dose of approximately 60 e^−^/Å^2^ and stored in electron event representation (EER) format. All image processing steps were carried out in cryoSPARC^54^. Movie frames were aligned and dose-weighted using patch-based motion correction^55,56^, followed by estimation of the contrast transfer function (CTF) using patch CTF estimation^56^.

For the apo dataset, 9879 micrographs were processed for particle picking, yielding approximately 2.0 million particles using a combination of blob picking and iterative Topaz-based particle picking^57^. Particles were extracted using a 480-pixel box size with 4x binning and subjected to several rounds of 2D classification to remove contaminants and poorly defined particles. The cleaned dataset was used for ab initio model generation, followed by heterogeneous refinement to separate well-resolved particles from junk classes. After iterative classification, a subset of 482,302 particles corresponding to the intact complex was selected. These particles were subjected to reference-based motion correction (Bayesian polishing) and subsequently re-extracted using a 480-pixel box size^58^. The polished particle stack was then refined using homogeneous and non-uniform refinement with C2 symmetry, resulting in a reconstruction at 2.33 Å resolution based on the gold-standard Fourier shell correlation (GSFSC = 0.143 criterion).

For the holo dataset, 16,650 micrographs were processed, yielding 892,879 particles after initial picking and classification. Ab initio reconstruction, followed by heterogeneous refinement and 3D classification, revealed multiple oligomeric states, which were separated into dimeric (~17.7%), trimeric (~50.6%), and tetrameric (~30.5%) populations. The particles belonging to each oligomeric class were subjected to reference-based motion correction (Bayesian polishing) and processed independently. The dimeric population was refined with C2 symmetry, yielding a reconstruction at 2.54 Å resolution. The trimeric and tetrameric assemblies were refined without imposed symmetry (C1), resulting in maps at 2.59 Å resolution. To further improve the map quality of flexible regions, focused local refinements were performed on individual protomeric units using soft masks. This yielded local reconstructions at 2.69 Å for the protomer within the trimer and 2.73 Å and 2.75 Å for the additional protomers within the tetrameric assembly, respectively. Subsequently, composite maps for the trimeric and tetrameric assemblies were generated in ChimeraX^59^ by first normalizing the individual maps to a common threshold using *vop scale*, followed by combining them with the *vop max* function. All maps were validated using gold-standard FSC, and local resolution variations were assessed. Both apo and holo maps were used for model building.

Initial models were generated using AlphaFold2^60^ and subsequently rigid-body fitted into the cryo-EM densities. Model building and manual adjustments were performed in ChimeraX and Coot^61,62^. Real-space refinement was carried out in PHENIX with appropriate stereochemical restraints. CIF restraints for Nickel (III) Ion (3NI) and Car-bonmonoxide-(dicyano) Iron (FCO) were generated using the eLBOW utility in PHENIX. Model quality was evaluated using standard validation metrics, including geometry statistics and map-to-model agreement. Final figures were prepared using UCSF ChimeraX.

### Sample preparation for the spectroscopic experiments

The DTT-reduced sample (as isolated) contained a 2 mM excess of the mild reductant DTT. The complete reduction of the HoxEFUYH protein was accomplished by adding an excess of 3 mM NaDT. For H_2_ reduction the HoxEFUYH protein solution was purged with a hydrated stream of pure H_2_ for one hour within an anaerobic glovebox containing 5% H_2_ and 95% N_2_. In order to study the effect of NAD^+^ binding, a tenfold excess of NAD^+^ was added to the respective sample. Ferredoxin reduction from H_2_ oxidation was investigated adjusting the HoxEFUYH to Fdx1 ratio to 1:2, while the reduction of HoxEFUYH by reduced Fdx1 was done at a ratio of 1:4 due to the partial reoxidation of the Fdx1 **(Extended Data Fig. 10e)**. To ensure, when investigating the effect from different sources of electrons (H_2_ or reduced Fdx1), that HoxEFUYH was completely oxidized, DTT was removed by consecutive dilution and concentration of the HoxEFUYH protein using 50 kDA Amicon® centrifugal filters. The respective samples were tested by EPR to confirm the full oxidation based on the absence of signals from reduced cofactors. Samples that were transported in gas tight vials during DTT removal steps were fully oxidized, likely due to the presence of residual O_2_ **(Extended Data Fig. 10a and 10e)**. On the other hand, samples that were kept in an anaerobic glove-box during the DTT removal stayed reduced and the EPR spectrum was identical to the DTT-reduced sample. To guarantee full oxidation of the later samples, they were purged with air for 5 min and EPR measured afterwards (Extended Data Fig. 10e). To prepare reduced NaDT-free Fdx1, this protein was first reduced with a 1 mM excess of NaDT. Subsequently, the chemical reductant was removed as described above using 10 kDA Amicon® centrifugal filters. During that process, a fraction of the protein reoxidized **(Extended data Fig. 10e)**.

### FTIR spectroscopy

The IR spectra displayed in Figs. 3d and 4e were measured in transmission mode using 0.03-0.1 mM solution of HoxEFUYH. The protein was loaded into a custom-made gastight IR transmission cell containing two CaF_2_ windows, which are separated by a 50 µm Teflon spacer. The sample compartment was continuously flushed with dry air, enabling observation of slow reoxidation **(Extended Data Fig. 10b)**, as O_2_ could slowly diffuse into the cell. Infrared spectra were collected on a Bruker Tensor 27 FTIR spectrometer equipped with a liquid nitrogen-cooled MCT detector at a spectral resolution of 2 cm^−1^. Due to the low concentration of the protein solution the corresponding second derivative spectra of the absorption data were calculated in order to clearly resolve the band positions unbiased from baseline corrections. Data processing and analysis were performed using Bruker OPUS software version 6.5 or higher.

All experiments shown in Extended Data Fig 10a were performed on hydrated protein films in attenuated total reflection (ATR) configuration using a FTIR spectrometer (Bruker Tensor27) equipped with a mercury cadmium telluride (MCT) detector cooled by liquid N_2_. All data were recorded with a spectral resolution of 2 cm^−1^ at 80 kHz scanning velocity and an average of 1.000 interferometer scans per spectrum. All experiments were conducted under the protective N_2_ atmosphere (pO_2_ < 2 ppm) of a Coy Lab anaerobic tent, at ambient temperature, ambient pressure, and in the dark. For each experiment, 1 µL of 0.03-0.01 mM HoxEFUYH protein solution was used forming a well-hydrated protein film as reported earlier^63^. The DTT-reduced protein and the DTT-free protein were first exposed to N_2_ for 30 min. Thereafter, reduction with pure H_2_ was done for 30 min. As the last step reoxidation was conducted for 10 min using 10% O_2_ and 90% N_2_. The composition of [NiFe]-hydrogenase cofactor intermediates was analyzed by calculating second derivative spectra within Bruker’s OPUS software version 7.0.

### EPR spectroscopy

The samples for EPR spectroscopy were measured at a concentration of 0.03-0.1 mM and a volume of 50-100 µl. The EPR spectra were recorded using a Bruker EMXplus spectrometer with an ER 4122 SHQE resonator (9.3 GHz) and an Oxford ESR900 helium-flow cryostat regulated by an Oxford ITC4 controller and a custom-built X-band spectrometer equipped with a Bruker SHQ resonator (9.4 GHz), an ESR 910 helium-flow cryostat, and an ITC503 temperature controller. The modulation amplitude was adjusted to 10 G and the modulation frequency to 100 kHz. Spectral simulations were carried out with the MATLAB-based EasySpin toolbox (version 6.0.12) employing the “pepper” function^64^. The Ni_a_-C-to-Ni_a_-L photoreaction was induced utilizing the focused light of a collimated 455-nm LED for illumination. The pure Ni_a_-L spectrum was generated by illuminating at 80 K for 2 hours. Subsequently, the temperature was increased to 200 K to purely enrich the Ni_a_-C state by thermal relaxation. To quantify the spin concentration of Fdx1 a mixture of 0.1 mM CuSO_4_ and 1 mM EDTA was used as external spin standard and measured under non-saturating conditions. The number of spins per mol for Fdx1 was determined to be 0.71.

## Data availability

The cryo-EM maps generated in this study have been deposited in the Electron Microscopy Data Bank (EMDB) under the following accession codes: EMD-57907 (HoxEFUYH_2_-apo), EMD-57893 (HoxEFUYH_2_-holo), EMD-57926 (composite map of HoxEFUYH_3_-holo), EMD-57914 (consensus map of HoxEFUYH_3_-holo), EMD-57925 (local refined map of the third protomer of HoxEFUYH_3_-holo), EMD-57948 (composite map of HoxEFUYH_4_-holo), EMD-57945 (consensus map of HoxEFUYH_4_-holo), EMD-57946 (local refined map of the third protomer of HoxEFUYH_4_-holo), EMD-57947 (local refined map of the fourth protomer of HoxEFUYH_4_-holo). Corresponding coordinate files have been deposited in the RCSB Protein Data Bank (PDB) under the following accession codes: 30PC (HoxEFUYH_2_-apo), 30OJ (HoxEFUYH_2_-holo), 30PR (HoxEFUYH_3_-holo), 30QE (HoxEFUYH_4_-holo). Structural and sequence data used for comparison with HoxEFUYH sub-units are available in the Protein Data Bank: 6Q8W (NADH:quinone oxidoreductase from *Thermus thermophilus*), 8A6T (electron bifurcating Fe-Fe hydrogenase HydABC complex from *Thermoanaerobacter kivui*), 5XF9 (soluble [NiFe]-hydrogenase HoxFUYH complex from *Thermus thermophilus*), 7T2R (electron bifurcating [NiFe]-hydrogenase HydABCSL complex from *Acetomicrobium mobile*). Strains and plasmids generated in this study are available from the corresponding authors upon request.

## Acknowledgements

We thank the cryo-EM Facility of Marburg University for technical support and the cryo-EM Platform at Helmholtz Munich, for cryo-EM data acquisition. We also thank Anuj Kumar for valuable suggestions during cryo-EM data processing and model building.

## Author contributions

J.M.S., J.A., C.L. and K.G. conceived and designed the study. G.P.M., N.S., M.B., C.L., S.T.S. and J.A. acquired the data. G.P.M., N.S., M.B., J.A., C.T., C.L., S.T.S., I.Z. and K.G. analyzed and interpreted the data. J.M.S., M.B., J.A., I.Z., and K.G. supervised the work. G.P.M., N.S., C.L., J.M.S., and K.G. drafted the manuscript. G.P.M., N.S., C.L., A.K., C.T., S.T.S., I.Z. J.A., J.M.S., and K.G. substantially revised the manuscript. J.M.S., I.Z. and K.G. acquired funding. All authors reviewed and approved the final version of the manuscript.

