## Extended Data and Supplementary Information for "Electron confurcation drives photosynthetic H_2_ production in cyanobacteria"

Extended and supplementary data:

---

**Extended Data Figures 1-10**
**Extended Data Tables 1-4**

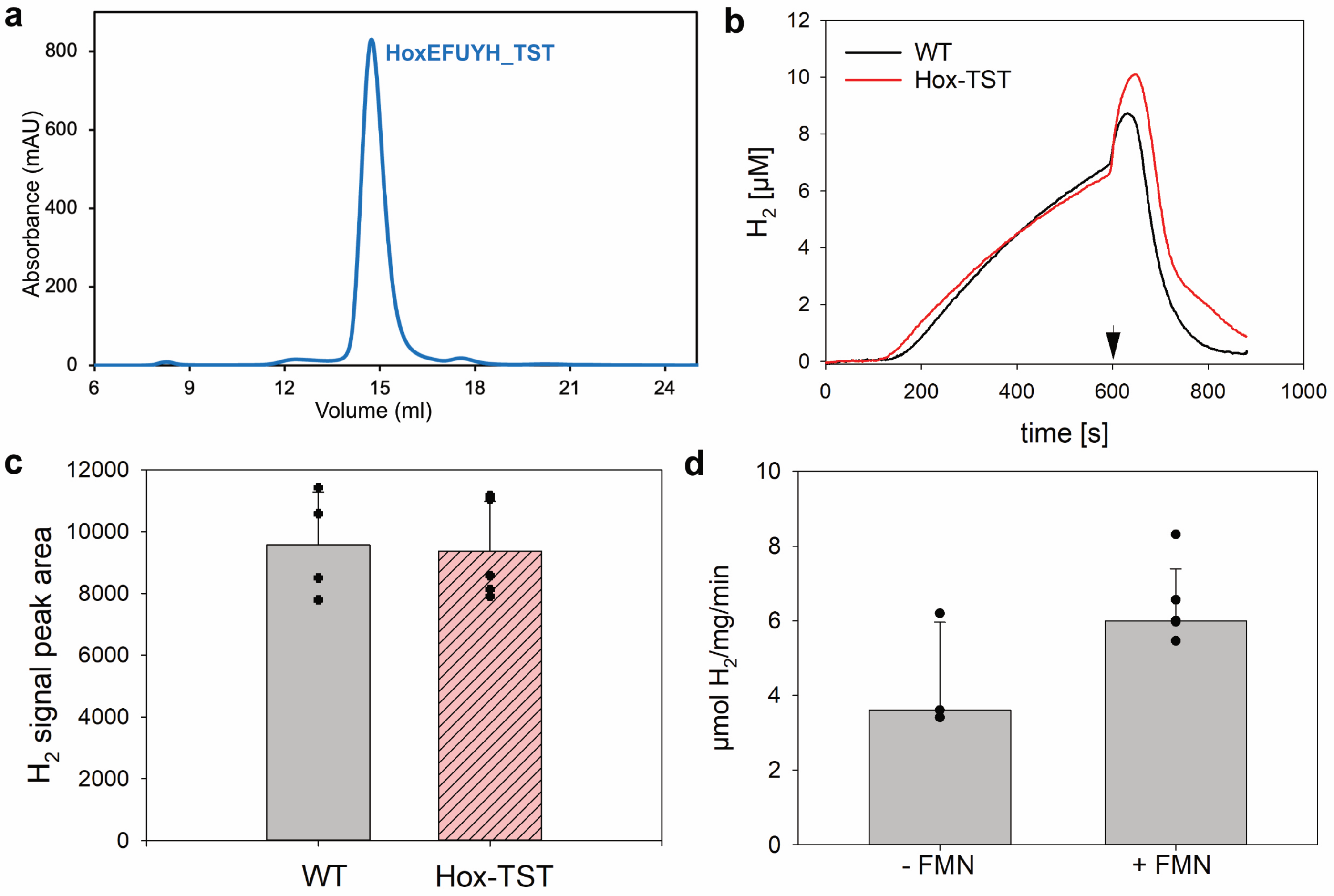

**Extended Data Fig. 1: Purification, in vivo activity, and cofactor dependence of cyanobacterial HoxEFUYH hydrogenase. a,** Representative SEC profile of anaerobically purified HoxEFUYH-TST from *Synechocystis* sp. PCC 6803. The major symmetric peak eluting at ~14.5 mL corresponds to the intact HoxEFUYH complex. **b,** Fermentative and photosynthetic $H_2$ production under anaerobic conditions (10 mM glucose, 4 U/$\mu$l glucose oxidase, 5 U/$\mu$l catalase) via Clark-type $H_2$-sensor (Unisense, Aarhus, Denmark). Cells were adapted to darkness for 10 minutes, followed by illumination (~800 $\mu$E, 625 nm) as indicated with an arrow. **c,** Methyl viologen assay via gas chromatography (Nexis™ GC-2030, Shimadzu, Kyoto, Japan). Cells were adjusted to $OD_{750}$ = 1.0 and incubated with 5 mM methyl viologen and 20 mM sodium dithionite for 30 minutes prior measurement. **d,** *In vitro* $H_2$ uptake rates in presence of $NAD^+$ and oxidized ferredoxin 1 without (n = 3) and with addition of 100 $\mu$M FMN (n = 5). $H_2$ concentration was recorded with a Clark-type $H_2$ sensor (Unisense, Aarhus, Denmark). Anaerobic conditions were achieved with an oxygen-scavenging enzyme mix (40 mM glucose, 16 U/$\mu$l glucose oxidase, 20 U/$\mu$l catalase).

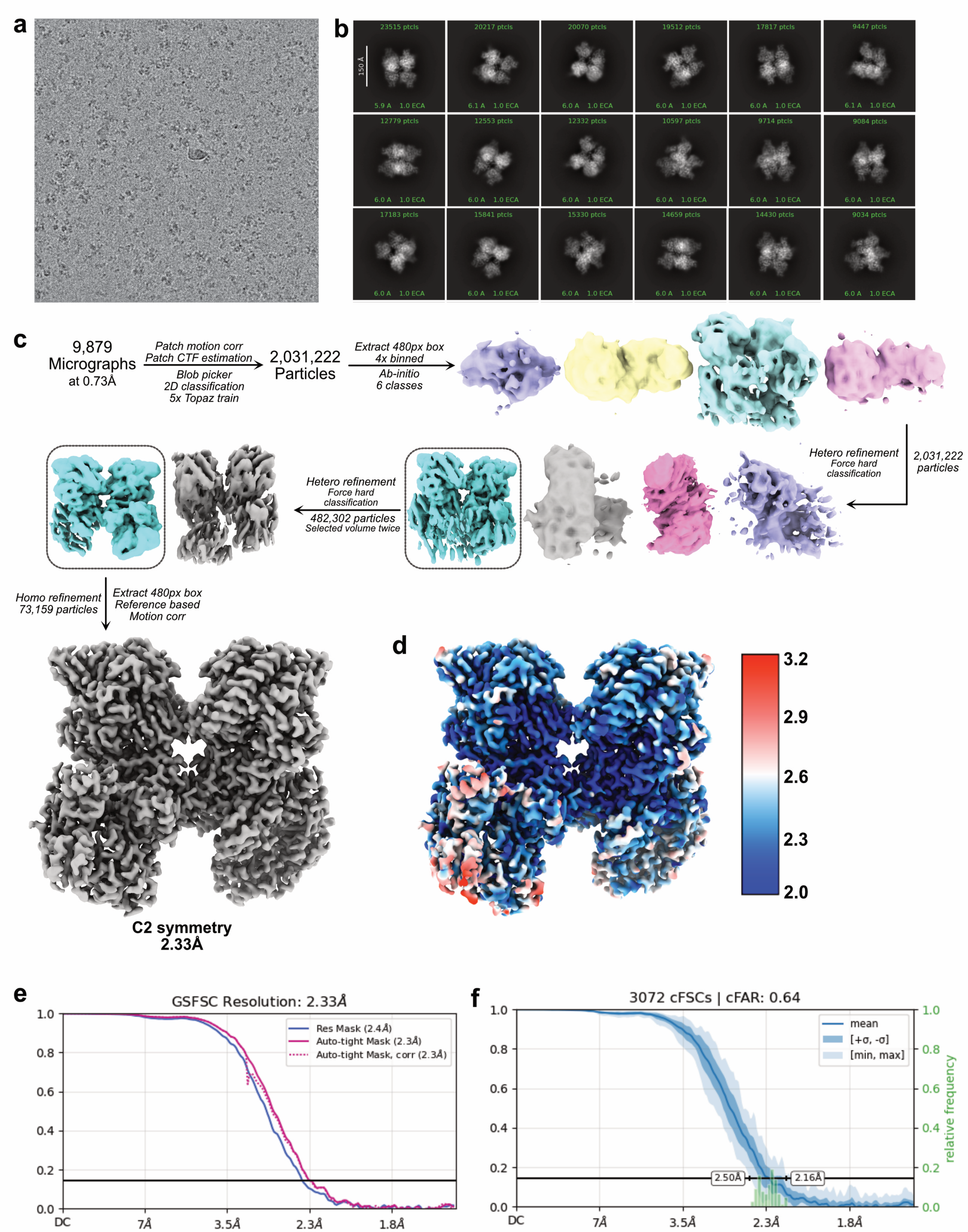

**Extended Data Fig. 2: Cryo-EM data processing and reconstruction of HoxEFUYH in the Apo state. a,** Representative cryo-EM micrograph of anaerobically purified HoxEFUYH. **b,** Reference-free 2D class averages showing different particle orientations. **c,** Cryo-EM image processing workflow for the apo HoxEFUYH dataset in cryoSPARC. **d,** Local resolution map of the final reconstruction colored according to local resolution estimates. **e,** GSFSC curves of the final reconstruction. **f,** Conical FSC (cFSC) analysis showing isotropic resolution distribution and limited orientation bias.

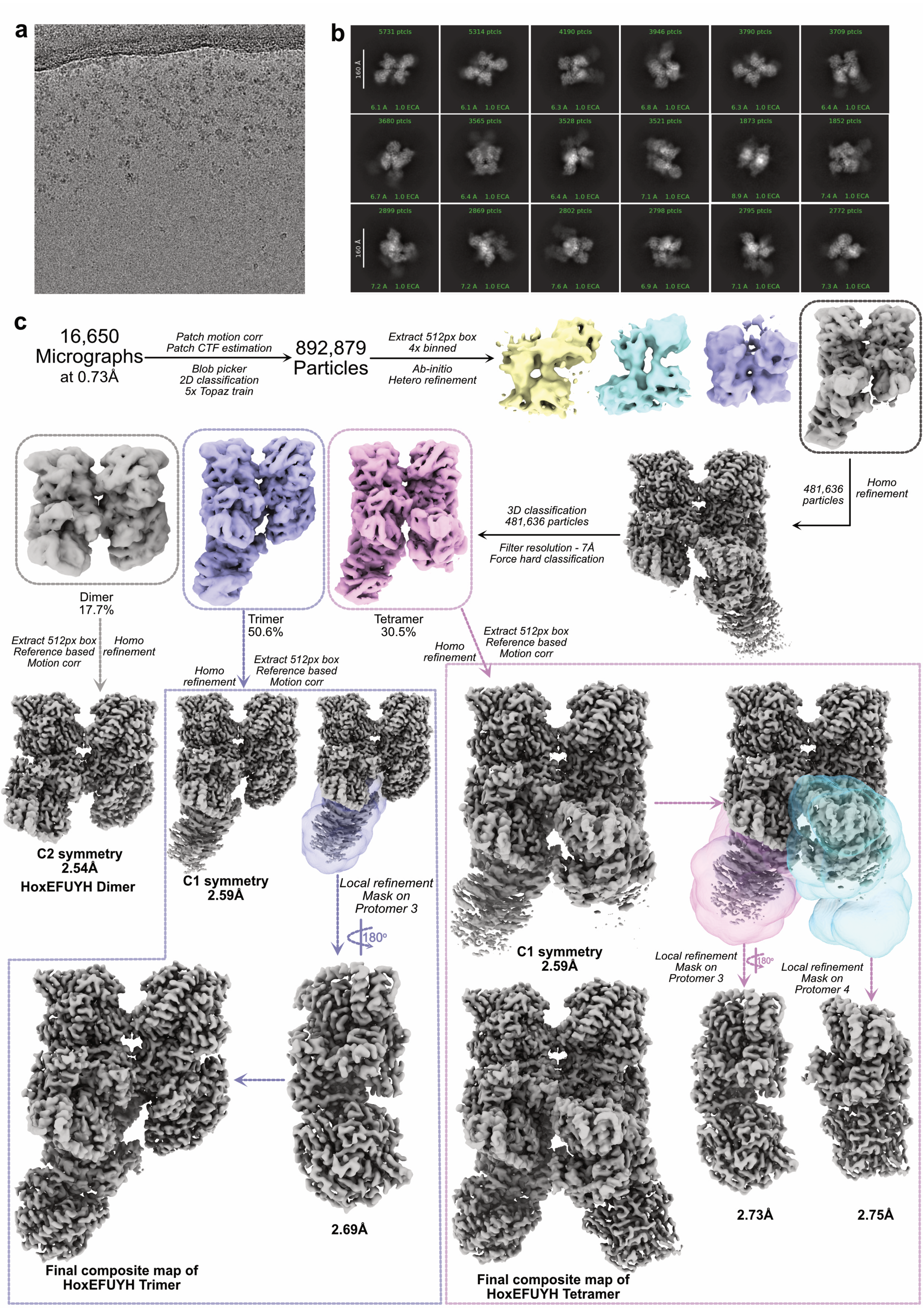
a
b
5731 ptcls
5314 ptcls
4190 ptcls
3946 ptcls
3790 ptcls
3709 ptcls
3680 ptcls
3565 ptcls
3528 ptcls
3521 ptcls
1873 ptcls
1852 ptcls
2899 ptcls
2869 ptcls
2802 ptcls
2798 ptcls
2795 ptcls
2772 ptcls
160 Å
c
16,650 Micrographs at 0.73Å
Patch motion corr
Patch CTF estimation
Blob picker
2D classification
5x Topaz train
892,879 Particles
Extract 512px box
4x binned
Ab-initio
Hetero refinement
481,636 particles
Homo refinement
3D classification
481,636 particles
Filter resolution - 7Å
Force hard classification
Dimer
17.7%
Trimer
50.6%
Tetramer
30.5%
Extract 512px box
Reference based
Motion corr
Homo refinement
C2 symmetry
2.54Å
HoxEFUYH Dimer
C1 symmetry
2.59Å
Local refinement
Mask on
Protomer 3
180°
2.69Å
Final composite map of
HoxEFUYH Trimer
C1 symmetry
2.59Å
Local refinement
Mask on
Protomer 3
Local refinement
Mask on
Protomer 4
2.73Å
2.75Å
Final composite map of
HoxEFUYH Tetramer

**Extended Data Fig. 3: Cryo-EM data processing of HoxEFUYH in the NADH bound state. a,** Representative cryo-EM micrograph of anaerobically purified HoxEFUYH. **b,** Reference-free 2D class averages showing different particle orientations. **c,** Cryo-EM image processing workflow for the apo HoxEFUYH dataset in cryoSPARC.

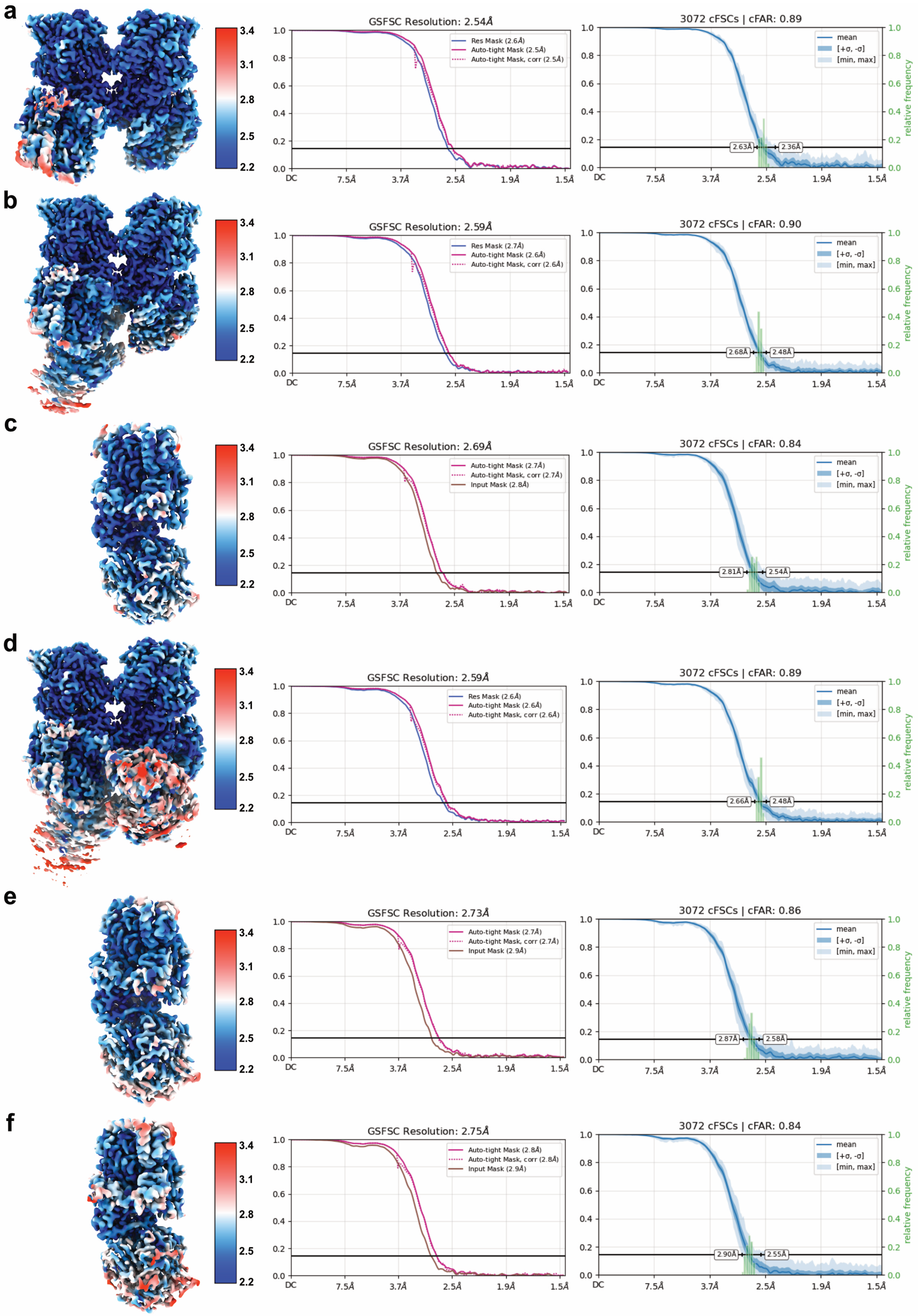

a
GSFSC Resolution: 2.54Å
Res Mask (2.6Å)
Auto-tight Mask (2.5Å)
Auto-tight Mask, corr (2.5Å)
3072 cFSCs | cFAR: 0.89
2.63Å
2.36Å
b
GSFSC Resolution: 2.59Å
Res Mask (2.7Å)
Auto-tight Mask (2.6Å)
Auto-tight Mask, corr (2.6Å)
3072 cFSCs | cFAR: 0.90
2.68Å
2.48Å
c
GSFSC Resolution: 2.69Å
Auto-tight Mask (2.7Å)
Auto-tight Mask, corr (2.7Å)
Input Mask (2.8Å)
3072 cFSCs | cFAR: 0.84
2.81Å
2.54Å
d
GSFSC Resolution: 2.59Å
Res Mask (2.6Å)
Auto-tight Mask (2.6Å)
Auto-tight Mask, corr (2.6Å)
3072 cFSCs | cFAR: 0.89
2.66Å
2.48Å
e
GSFSC Resolution: 2.73Å
Auto-tight Mask (2.7Å)
Auto-tight Mask, corr (2.7Å)
Input Mask (2.9Å)
3072 cFSCs | cFAR: 0.86
2.87Å
2.58Å
f
GSFSC Resolution: 2.75Å
Auto-tight Mask (2.8Å)
Auto-tight Mask, corr (2.8Å)
Input Mask (2.9Å)
3072 cFSCs | cFAR: 0.84
2.90Å
2.55Å
mean
[+σ, -σ]
[min, max]
relative frequency
DC
7.5Å
3.7Å
2.5Å
1.9Å
1.5Å
3.4
3.1
2.8
2.5
2.2

**Extended Data Fig. 4: Cryo-EM map reconstruction and resolution analysis of HoxEFUYH in the NADH-bound state. a,** Local resolution map, GSFSC, and conical FSC (cFSC) analysis of the dimeric HoxEFUYH reconstruction. **b,** Local resolution map, GSFSC, and cFSC analysis of the trimeric HoxEFUYH reconstruction. **c,** Local resolution map, GSFSC, and cFSC analysis of the local refined protomer 3 from the trimeric HoxEFUYH assembly. **d,** Local resolution map, GSFSC, and cFSC analysis of the tetrameric HoxEFUYH reconstruction. **e,** Local resolution map, GSFSC, and cFSC analysis of the local refined protomer 3 from the tetrameric HoxEFUYH assembly. **f,** Local resolution map, GSFSC, and cFSC analysis of the local refined protomer 4 from the tetrameric HoxEFUYH assembly.

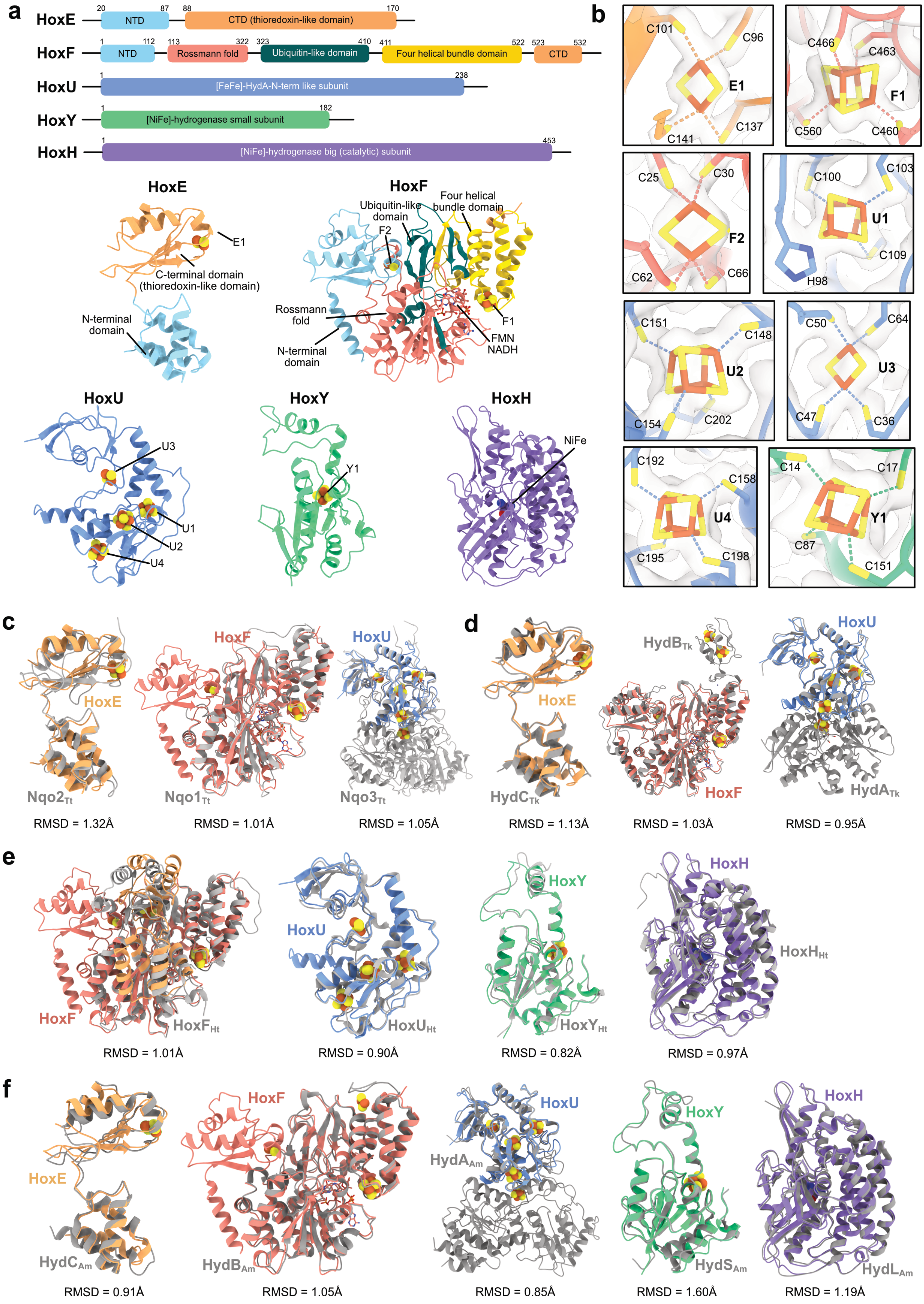

a
HoxE
20 NTD 87
88 CTD (thioredoxin-like domain) 170
HoxF
1 NTD 112
113 Rossmann fold 322
323 Ubiquitin-like domain 410
411 Four helical bundle domain 522
523 CTD 532
HoxU
1 [FeFe]-HydA-N-term like subunit 238
HoxY
1 [NiFe]-hydrogenase small subunit 182
HoxH
1 [NiFe]-hydrogenase big (catalytic) subunit 453
HoxE
E1
C-terminal domain (thioredoxin-like domain)
N-terminal domain
HoxF
Ubiquitin-like domain
Four helical bundle domain
F2
Rossmann fold
N-terminal domain
F1
FMN
NADH
HoxU
U3
U1
U2
U4
HoxY
Y1
HoxH
NiFe
b
C101
C96
E1
C141
C137
C466
C463
F1
C560
C460
C25
C30
F2
C62
C66
C100
C103
U1
C109
H98
C151
C148
U2
C154
C202
C50
C64
U3
C47
C36
C192
C158
U4
C195
C198
C14
C17
Y1
C87
C151
c
HoxE
Nqo2$_{Tt}$
RMSD = 1.32Å
HoxF
Nqo1$_{Tt}$
RMSD = 1.01Å
HoxU
Nqo3$_{Tt}$
RMSD = 1.05Å
d
HoxE
HydC$_{Tk}$
RMSD = 1.13Å
HydB$_{Tk}$
HoxF
RMSD = 1.03Å
HoxU
HydA$_{Tk}$
RMSD = 0.95Å
e
HoxF
HoxF$_{Ht}$
RMSD = 1.01Å
HoxU
HoxU$_{Ht}$
RMSD = 0.90Å
HoxY
HoxY$_{Ht}$
RMSD = 0.82Å
HoxH
HoxH$_{Ht}$
RMSD = 0.97Å
f
HoxE
HydC$_{Am}$
RMSD = 0.91Å
HoxF
HydB$_{Am}$
RMSD = 1.05Å
HoxU
HydA$_{Am}$
RMSD = 0.85Å
HoxY
HydS$_{Am}$
RMSD = 1.60Å
HoxH
HydL$_{Am}$
RMSD = 1.19Å

**Extended Data Fig. 5: Domain organization, cofactor coordination, and structural comparison of cyanobacterial HoxEFUYH subunits. a,** Domain architecture and cryo-EM structures of the HoxEFUYH subunits. Top, schematic representation of domain organization in HoxE, HoxF, HoxU, HoxY, and HoxH. Bottom, experimentally determined cryo-EM structures highlighting FeS clusters, FMN, NADH, and the catalytic NiFe center. **b,** Electron-density maps of the FeS cluster coordination environments in HoxEFUYH. Close-up views of the E1, F1, F2, U1, U2, U3, U4, and Y1 clusters with coordinating residues. **c,** Structural comparison of HoxE, HoxF, and HoxU with the homologous complex I subunits Nqo2, Nqo1, and Nqo3 from *Thermus thermophilus* (PDB: 6Q8W). HoxE aligns closely with Nqo2, whereas HoxF contains an additional N-terminal domain absent in Nqo1. HoxU lacks the C-terminal domain present in Nqo3. RMSD values are indicated below each comparison. **d,** Structural comparison of HoxE, HoxF, and HoxU with the corresponding HydC, HydB, and HydA subunits from the bifurcating hydrogenase of *Thermotoga kivui* (PDB: 8A6T). HoxE closely aligns with HydC, whereas HoxF lacks the flexible C-terminal ferredoxin-like extension present in HydB. HoxU aligns with the N-terminal FeS-containing region of HydA but lacks the large C-terminal domain extension observed in HydA. RMSD values are indicated below each comparison. **e,** Structural comparison of HoxF, HoxU, HoxY, and HoxH with homologous subunits from *Hydrogenophilus thermoluteolus* (PDB: 5XF9). HoxF and HoxU closely align with their homologs, whereas minor differences are observed in peripheral loops and terminal regions. The catalytic hydrogenase subunits HoxY and HoxH are highly conserved. RMSD values are indicated below each comparison. **f,** Structural comparison of HoxE, HoxF, HoxU, HoxY, and HoxH with homologous subunits from the bifurcating hydrogenase of *Acetomicrobium mobile* (PDB: 7T2R). HoxE and HoxF closely align with HydC and HydB, respectively, whereas HoxU lacks the extended C-terminal region present in HydA that contains the active site H-cluster of FeFe-hydrogenases. HoxY and HoxH show overall conservation of the hydrogenase small and catalytic subunits. RMSD values are indicated below each comparison.

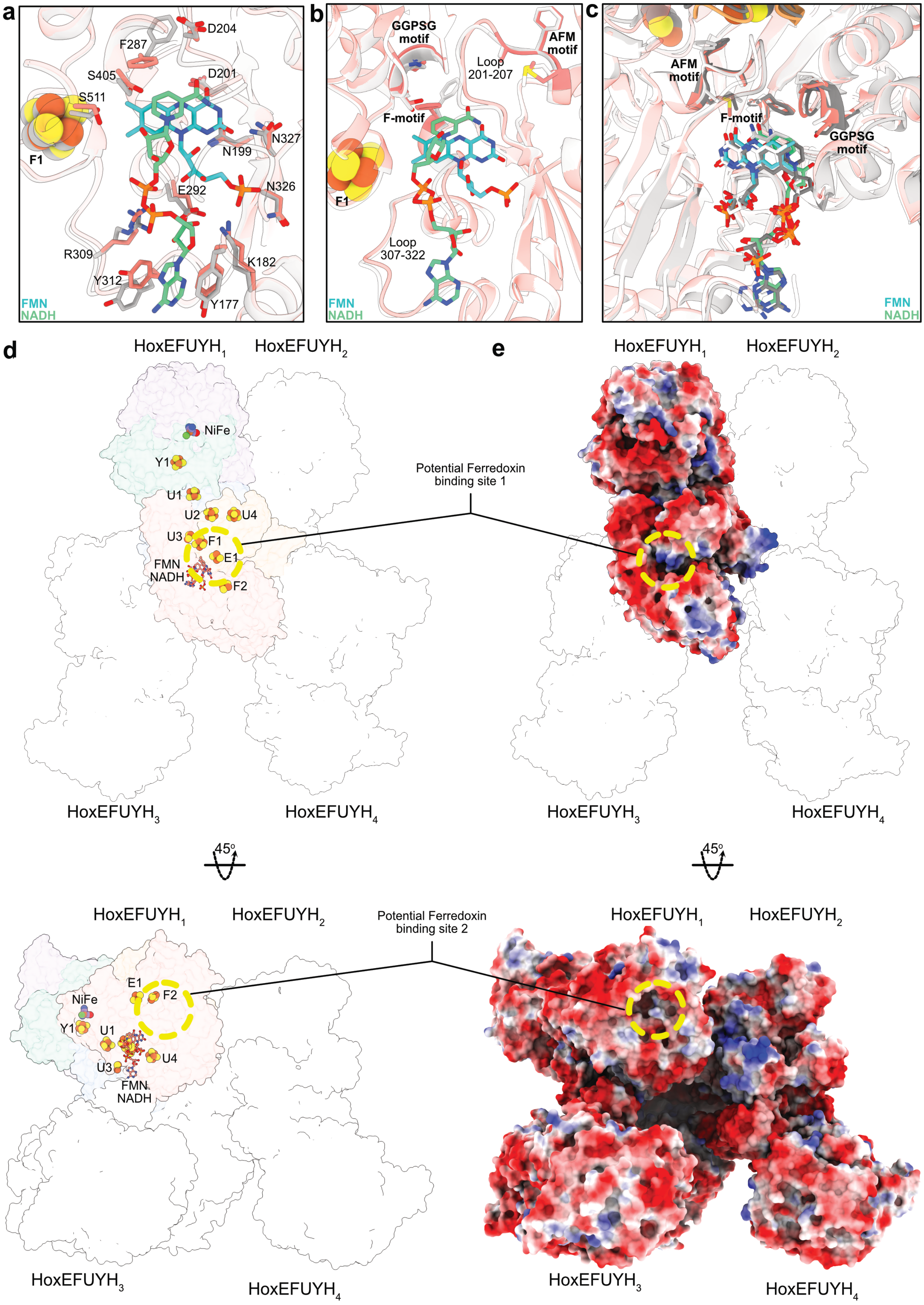

a
D204
F287
S405
D201
S511
N327
N199
F1
N326
E292
R309
K182
Y312
Y177
FMN
NADH
b
GGPSG motif
AFM motif
Loop 201-207
F-motif
F1
Loop 307-322
FMN
NADH
c
AFM motif
F-motif
GGPSG motif
FMN
NADH
d
HoxEFUYH$_1$
HoxEFUYH$_2$
NiFe
Y1
U1
U2
U4
U3
F1
E1
FMN
NADH
F2
HoxEFUYH$_3$
HoxEFUYH$_4$
e
HoxEFUYH$_1$
HoxEFUYH$_2$
Potential Ferredoxin binding site 1
HoxEFUYH$_3$
HoxEFUYH$_4$
45º
45º
HoxEFUYH$_1$
HoxEFUYH$_2$
Potential Ferredoxin binding site 2
E1
F2
NiFe
Y1
U1
U4
U3
FMN
NADH
HoxEFUYH$_3$
HoxEFUYH$_4$
HoxEFUYH$_1$
HoxEFUYH$_2$
HoxEFUYH$_3$
HoxEFUYH$_4$

**Extended Data Fig. 6: FMN/NADH-binding architecture, conserved bifurcation motifs, and potential ferredoxin-binding sites in HoxEFUYH.** **a,** FMN–NADH binding site in HoxF comparing the apo (silver) and NADH-bound (salmon) structures. Close-up view of the FMN/NADH-binding pocket highlighting residues involved in cofactor coordination and local structural rearrangements upon NADH binding. **b,** Structural organization of the FMN/NADH-binding region highlighting the conserved AFM, F-, and GGPSG motifs characteristic of bifurcating hydrogenases. Comparison between the apo (silver) and NADH-bound (salmon) structures reveals local loop rearrangements surrounding the FMN/NADH-binding pocket upon NADH binding. **c,** Structural comparison of the FMN/NADH-binding region between HoxEFUYH-NADH (salmon), HydABC-NADH-bound (tin), and HydABCSL-NADH-bound (snow) structures. The overall FMN/NADH-binding architecture and the conserved AFM, F-, and GGPSG motifs are structurally conserved across the complexes. **d-e,** Putative ferredoxin-binding regions in HoxEFUYH shown as cartoon and electrostatic surface representations. Positively charged surface patches adjacent to exposed E1/F2 FeS clusters of the low-potential branch may facilitate transient interaction with negatively charged ferredoxin. Electrostatic surface potentials are colored from red (negative) to blue (positive).

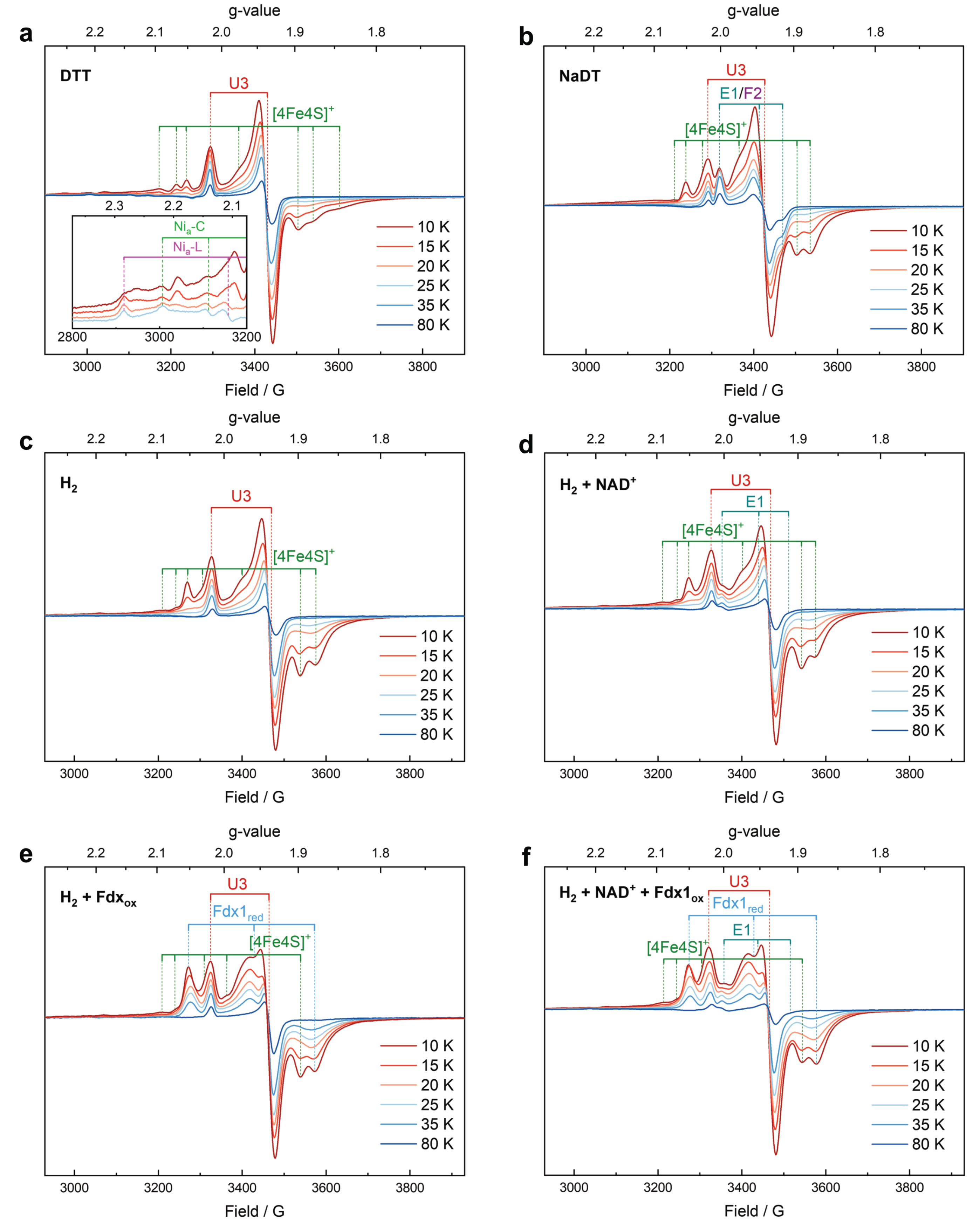

**Extended Data Fig. 7: Temperature dependent EPR spectra of HoxEFUYH recorded at 1 mW microwave power. a,** DTT-reduced. The inset shows a magnification of the spectral region characteristic for signals from the [NiFe] site. The EPR signals of $Ni_a$-C and $Ni_a$-L exhibit line broadening and partial splitting, indicative of magnetic coupling to a nearby paramagnetic center. Based on the structural arrangement, the proximal cluster Y1 is the only plausible candidate, suggesting that this [4Fe-4S] cluster is one-electron reduced under these mildly reducing conditions. In addition, the characteristic axial signal of the reduced [2Fe2S] cluster U3 is observed, consistent with its previously determined midpoint potential of –349 mV[1]. Upon decreasing the temperature to 10 K, up to four additional EPR signals emerge (g = 2.09, 2.07, 2.05, 2.03, 1.99, 1.97, 1.90, 1.88. 1.84)

attributable to up to four reduced [4Fe-4S] clusters[2]. However, strong spectral overlap precludes unambiguous assignment of the individual cofactors. **b,** NaDT-reduced. **c,** $H_2$-reduced. **d,** $H_2$-reduced in the presence of $NAD^+$. **e,** $H_2$-reduced in the presence of oxidized Fdx1. **f,** $H_2$-reduced in the presence of $NAD^+$ and oxidized Fdx1.

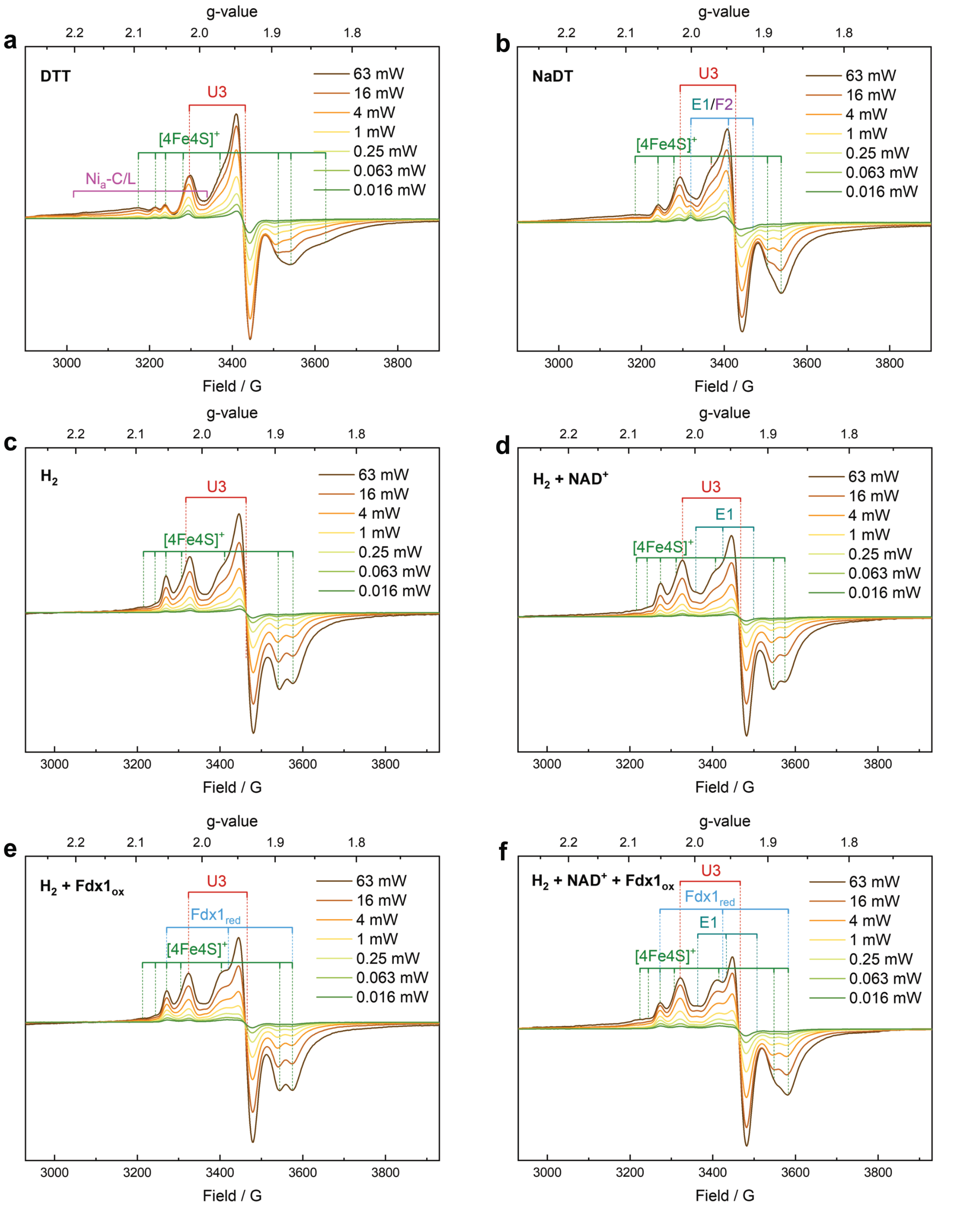

**Extended Data Fig. 8: Power dependent EPR spectra of HoxEFUYH recorded at 10 K. a,** DTT-reduced. **b,** NaDT-reduced. **c,** $H_2$-reduced. **d,** $H_2$-reduced in the presence of $NAD^+$. **e,** $H_2$-reduced in the presence of oxidized Fdx1. **f,** $H_2$-reduced in the presence of $NAD^+$ and oxidized Fdx1.

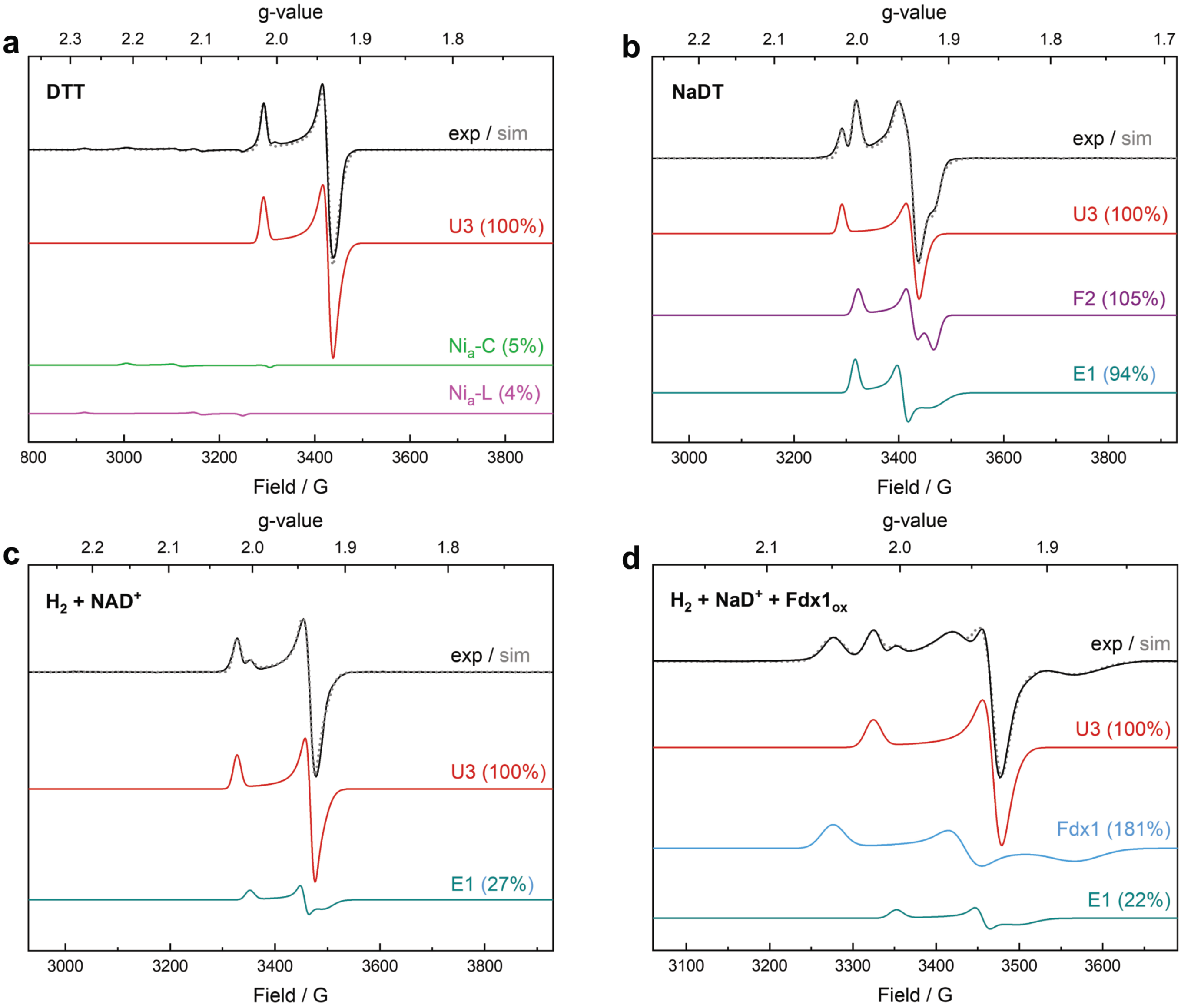

**Extended Data Fig. 9: Numerical simulation of the EPR spectra of HoxEFUYH recorded at 35 K and 1 mW.** The signal from the [2Fe-2S] cluster U3 has the highest reduction potential (-349 mV)[1] and was used as an internal reference for normalizing the individual spectra. **a,** DTT-reduced. **b,** NaDT-reduced. **c,** $H_2$-reduced in the presence of $NAD^+$. **d,** $H_2$-reduced in the presence of $NAD^+$ and oxidized Fdx1.

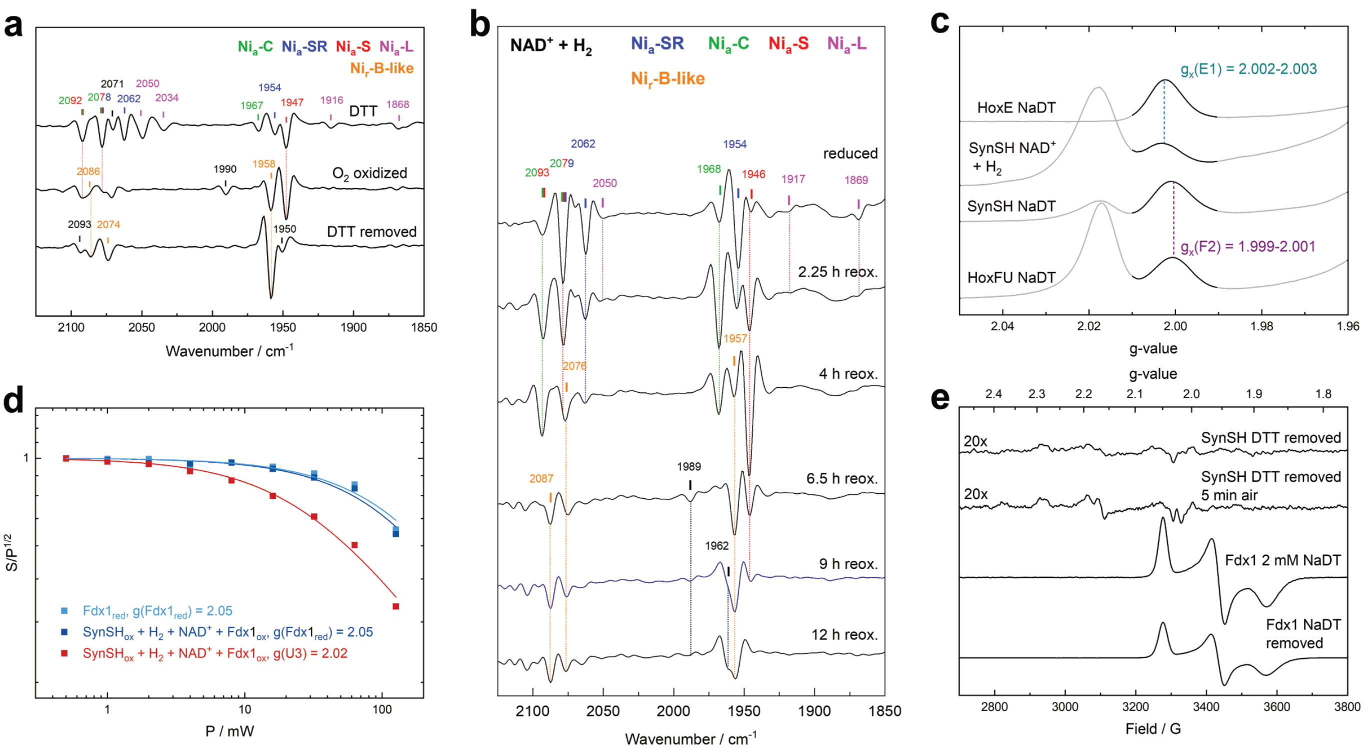

**Extended Data Fig. 10: IR and EPR spectroscopic data of HoxEFUYH and related proteins. a,** ATR-IR spectra of hydrated protein films of HoxEFUYH recorded at 293 K, reduced with DTT and incubated under a $N_2$-atmosphere (top), oxidized with a gas mix (10% $O_2$ and 90% $N_2$ ) after $H_2$-reduction (middle) and incubated with $N_2$ after removal of DTT (bottom). While the first spectrum exhibits a mixture of catalytic active states in line with the transmission IR data (Fig. 3D) incubation with $O_2$ induces the oxidation of the [NiFe] center, forming the Ni$_r$-B-like state and the most oxidized catalytic state Ni$_a$-S, consistent with the IR data recorded in solution (Fig. 4e)[3]. The removal of DTT in the presence of trace oxygen (see Methods) caused the almost pure enrichment of the oxidized, but EPR-silent Ni$_r$-B like state[3]. **b,** IR spectra of HoxEFUYH solution recorded at 283 K in transmission mode. After reducing the protein with pure $H_2$ the enzyme slowly reoxidizes due to diffusion of air into the transmission cell. During the reoxidation the sequential enrichment of the Ni$_a$-SR, Ni$_a$-C, Ni$_a$-S and Ni$_r$-B-like state was observed[3]. **c,** Magnification of the spectral region characteristic for the $g_x$ value of the EPR signal from E1 and F2 [2Fe-2S] clusters from EPR spectra of HoxEFUYH and isolated subunits recorded at 35 K and 1 mW microwave power. EPR spectrum of the isolated HoxE subunit reduced with an excess of NaDT, containing only E1 (top)[4], HoxEFUYH $H_2$-reduced in the presence of NAD$^+$ with E1 partially reduced (second from the top), HoxEFUYH reduced with an excess of NaDT with E1 and F2 completely reduced (second from the bottom), isolated HoxFU subunit reduced with an excess of NaDT exhibiting signals from reduced F2 and U3 (bottom)[4]. **d,** Power saturation data of the signal from isolated reduced Fdx1 as well as Fdx1 mixed with HoxEFUYH and incubated with $H_2$ in the presence of NAD$^+$ and the signal from U3, respectively measured at $g_x$ position and fitted to the empirical equation from Portis and Castner[5]. **e,** EPR spectra of HoxEFUYH and Fdx1 recorded at 35 K and 1 mW microwave power. EPR spectrum of HoxEFUYH after removal of DTT by consecutive diluting and concentrating steps, which was slowly oxidized during transport by residual oxygen (top). A very weak signal from a paramagnetic Ni species ($g_x$ = 2.28, $g_y$ = 2.17, $g_z$ = 2.03) was observed. A second weak Ni species ($g_x$ = 2.38, $g_y$ = 2.18, $g_z$ = 2.02) could be detected when the enzyme was incubated with air for 5 minutes directly after removing the DTT (second from the top, see also Methods). EPR spectra of isolated Fdx1 reduced with an excess of NaDT (second from the bottom) and after removal of the NaDT (bottom). During the consecutive diluting and concentrating steps 40% of Fdx1 was oxidized, possibly due to unspecific side reactions, which explains the observed loss of signal intensity.

| Primer name | Sequence (5'-3') | Amplified fragment | Generated plasmid |
|---|---|---|---|
| hoxout1-tar | CGAAGCAGGGTTATGCAGCG-GAAAGTATACCTTAAC-CGCCGTTTTTATCTGCCAGTGAAGCCCTT | upstream re-combination site | pMQ80hox (backbone) |
| hoxin2-tar | TAGA-TATGCCCTCGGTGATGCCATTTAAATGA-TAAAAGATGATTGGGAGA | | |
| hoxin1-tar | TCTCCCAATCATCTTTTATCATTTAAATGG-CATCACCGAGGGCATATCTA | downstream re-combination site | |
| hoxout2-tar | ATTTGCACGGCGTCACACTTTGCTATGCCA-TAGCATTTTTATCCATAA-GATTAGCGTCCCCACGGCACTGGCACTCTT | | |
| hox1-psbA_40 | TTTAC-CTGGTTTAGGCTCTCCCAATCATCTTTTATCATTTGATTGCGGCTTTAGCGTTCCA | PsbAII-promo-tor + *hoxE* and *hoxF* | pP$_{PsbAII}$*Syn*-Hox-TST |
| hoxU-hoxF_2 | AACAACAGACATT-GGCCAAAAATCTCCTGAAAATGAACTA-GACTTTGAGTAATTCTTCATA | | |
| hoxF-hoxU_2 | AAGAATTACTCAAAGTCTAG-TTCATTTTCAGGAGATTTTT-GGCCAATGTCTGTT | *hoxU* | |
| hoxY-hoxU | GCCATGATTAAAATCTCCTAGACTAC-CTGACCCATTCCTTTTCT | | |
| hoxU-hoxY | AGAAAAGGAATGGGTCAGGTAGTCTAGGA-GATTTTAATCATGGCTA | *hoxY* and *hoxH* with N-termi-nal Twin-Strep tag | |
| Sp-hoxH_Syn | CACCCAGCCTGCGCGATTAATCCCGCTG-GATGGACTTAATTA | | |
| hoxH-Sp_Syn | AGTCCATCCAGCGGGATTAATCGCG-CAGGCTGGGTG | Spectinomycin resistance | |
| hox2-Sp_40 | GAGTTCTCCTAGA-TATGCCCTCGGTGATGCCGGTTGATTTTT-GATTTGCCCTCGCTAGATTTTA | | |

**Supplementary Table 1: List of primers used for cloning of pP$_{PsbAII}$*Syn*Hox-TST and respective backbone (pMQ80hox).**

| | HoxEFUYH-Apo dimer (EMDB-57907) (PDB 30PC) | HoxEFUYH-NADH bound dimer (EMDB-57893) (PDB 30OJ) | HoxEFUYH-NADH bound trimer (Composite map and model) (EMDB-57926) (PDB 30PR) | HoxEFUYH-NADH bound tri-mer (Third Protomer Local Refinement) (EMDB-57925) |
|---|---|---|---|---|
| **Data collection and processing** | | | | |
| Magnification | 165,000x | 165,000x | 165,000x | 165,000x |
| Voltage (kV) | 300 | 300 | 300 | 300 |
| Electron exposure (e–/Å$^2$) | 60 | 60 | 60 | 60 |
| Defocus range (μm) | 0.5-2.0 | 0.5-2.0 | 0.5-2.0 | 0.5-2.0 |
| Pixel size (Å) | 0.73 | 0.73 | 0.73 | 0.73 |
| Symmetry imposed | C2 | C2 | C1 | C1 |
| Initial particle images (no.) | 2,031,222 | 892,879 | 892,879 | 892,879 |
| Final particle images (no.) | 73,159 | 82,841 | 234,672 | 234,672 |
| Map resolution (Å) | 2.33 | 2.54 | 2.59 | 2.69 |
| FSC threshold | 0.143 | 0.143 | 0.143 | 0.143 |
| Map resolution range (Å) | 2.0-3.2 | 2.2-3.4 | 2.2-3.4 | 2.2-3.4 |
| **Refinement** | | | | |
| Initial model used (PDB code) | *de novo* (AlphaFold) | *de novo* (AlphaFold) | *de novo* (AlphaFold) | - |
| Model resolution (Å) | 2.33 | 2.54 | 2.59 | - |
| FSC threshold | 0.143 | 0.143 | 0.143 | - |
| Model resolution range (Å) | 2.3-2.7 | 2.3-2.9 | 1.8-3.0 | - |
| Map sharpening *B* factor (Å$^2$) | 57.8 | 76.7 | 81.5 | 85.5 |
| Model composition | | | | |
| Non-hydrogen atoms | 23964 | 24122 | 35966 | - |
| Protein residues | 3098 | 3099 | 4623 | - |
| Ligands | SF4: 10, 3NI: 2, FES: 6, FCO: 2 | FMN: 2, NAD: 2, SF4: 10, 3NI: 2, FES: 6, FCO: 2 | FMN: 3, NAD: 3, SF4: 15, 3NI: 3, FES: 9, FCO: 3 | - |
| *B* factors (Å$^2$) | | | | |
| Protein | 95.52 | 83.80 | 98.26 | - |
| Ligand | 78.98 | 80.72 | 85.48 | - |
| R.m.s. deviations | | | | |
| Bond lengths (Å) | 0.004 | 0.002 | 0.003 | - |
| Bond angles (°) | 0.584 | 0.503 | 0.540 | - |
| Validation | | | | |
| MolProbity score | 1.38 | 1.33 | 1.45 | - |
| Clashscore | 4.38 | 4.84 | 5.32 | - |
| Poor rotamers (%) | 1.36 | 0.86 | 1.25 | - |
| Ramachandran plot | | | | |
| Favored (%) | 97.73 | 97.63 | 97.54 | - |
| Allowed (%) | 2.27 | 2.37 | 2.46 | - |
| Disallowed (%) | 0.00 | 0.00 | 0.00 | - |

| | HoxEFUYH-NADH bound tetramer (EMDB-57948) (PDB 30QE) | HoxEFUYH-NADH bound tetramer (Third Protomer Local Refinement) (EMDB-57946) | HoxEFUYH-NADH bound tetramer (Fourth Protomer Local Refinement) (EMDB-57947) |
|---|---|---|---|
| **Data collection and processing** | | | |
| Magnification | 165,000x | 165,000x | 165,000x |
| Voltage (kV) | 300 | 300 | 300 |
| Electron exposure (e–/$Å^2$) | 60 | 60 | 60 |
| Defocus range (μm) | 0.5-2.0 | 0.5-2.0 | 0.5-2.0 |
| Pixel size (Å) | 0.73 | 0.73 | 0.73 |
| Symmetry imposed | C1 | C1 | C1 |
| Initial particle images (no.) | 892,879 | 892,879 | 892,879 |
| Final particle images (no.) | 139,319 | 139,319 | 139,319 |
| Map resolution (Å) | 2.59 | 2.73 | 2.75 |
| FSC threshold | 0.143 | 0.143 | 0.143 |
| Map resolution range (Å) | 2.2-3.4 | 2.2-3.4 | 2.2-3.4 |
| **Refinement** | | | |
| Initial model used (PDB code) | *de novo* (AlphaFold) | - | - |
| Model resolution (Å) | 2.59 | - | - |
| FSC threshold | 0.143 | - | - |
| Model resolution range (Å) | 1.7-3.0 | - | - |
| Map sharpening *B* factor ($Å^2$) | 72.5 | 75.4 | 77.9 |
| Model composition | | | |
| Non-hydrogen atoms | 47810 | - | - |
| Protein residues | 6147 | - | - |
| Ligands | FMN: 4, NAD: 4, SF4: 20, 3NI: 4, FES: 12, FCO: 4 | - | - |
| *B* factors ($Å^2$) | | | |
| Protein | 104.32 | - | - |
| Ligand | 85.63 | - | - |
| R.m.s. deviations | | | |
| Bond lengths (Å) | 0.003 | - | - |
| Bond angles (°) | 0.557 | - | - |
| Validation | | | |
| MolProbity score | 1.51 | - | - |
| Clashscore | 5.57 | - | - |
| Poor rotamers (%) | 1.32 | - | - |
| Ramachandran plot | | | |
| Favored (%) | 97.41 | - | - |
| Allowed (%) | 2.59 | - | - |
| Disallowed (%) | 0.00 | - | - |

**Supplementary Table 2: Cryo-EM data collection, refinement and validation statistics**

| Redox State | ν (CO) / $cm^{-1}$ | ν $(CN)_{as}$ / $cm^{-1}$ | ν $(CN)_s$ / $cm^{-1}$ | Reference |
|---|---|---|---|---|
| $Ni_r$-B-like | 1957 | 2076 | 2087 | This work |
| | 1957 | 2076 | 2088 | Germer et al[3]. |
| $Ni_a$-S | 1946 | 2077 | 2093 | This work |
| | 1947 | 2078 | 2093 | Germer et al[3]. |
| $Ni_a$-L | 1917<br>1869* | 2034 | 2050 | This work |
| $Ni_a$-C | 1968 | 2079 | 2093 | This work |
| | 1968 | 2079 | 2093 | Germer et al[3]. |
| $Ni_a$-SR | 1954 | 2062 | 2079 | This work |
| | 1955 | 2063 | 2079 | Germer et al[3]. |

*The two CO vibration are related to two separate $Ni_a$-L states, which most likely differ in the protonation state of the active site[6].

**Supplementary Table 3: Characteristic bands related to CO and CN stretching vibrations in the redox states of HoxEFUYH in comparison to published data.**

| FeS cluster / [NiFe] Redox State | Protein | $g_x$ | $g_y$ | $g_z$ | Reference |
|---|---|---|---|---|---|
| U3 | HoxEFUYH | 2.017 | 1.939 | 1.931 | This work |
| | HoxEFU | 2.017 | 1.939 | 1.929 | Lettau et al[4]. |
| | HoxEFU | 2.017 | 1.940 | 1.931 | Blahut et al[1]. |
| | HoxU | 2.017 | 1.944 | 1.931 | Blahut et al[1]. |
| E1 | HoxEFUYH | 2.002 | 1.948 | 1.915 | This work |
| | HoxEFU | 2.002 | 1.945 | 1.915 | Lettau et al[4]. |
| | HoxEFU | 2.003 | 1.945 | 1.915 | Blahut et al[1]. |
| | HoxE | 2.002 | 1.945 | 1.915 | Lettau et al[4]. |
| | HoxE | 2.003 | 1.945 | 1.915 | Blahut et al[1]. |
| F2 | HoxEFUYH | 1.999 | 1.939 | 1.915 | This work |
| | HoxEFU | 1.999 | 1.937 | 1.916 | Blahut et al[1]. |
| | HoxFU | 2.001 | 1.945 | 1.915 | Lettau et al[4]. |
| Fdx1 | Fdx1 | 2.049 | 1.955 | 1.880 | This work |
| | Fdx1 | 2.048 | 1.955 | 1.877 | Boehm et al[7]. |
| $Ni_a$-L | HoxEFUYH | 2.275 | 2.105 | 2.043 | This work |
| $Ni_a$-C | HoxEFUYH | 2.210 | 2.135 | 2.011 | This work |

**Supplementary Table 4: Observed and previously reported g-values of the [2Fe-2S] clusters and paramagnetic [NiFe] redox states in Hox-EFUYH and Fdx1.**
